# Structure-function analysis of the cyclic β-1,2-glucan synthase

**DOI:** 10.1101/2023.05.05.539553

**Authors:** Jaroslaw Sedzicki, Dongchun Ni, Frank Lehmann, Henning Stahlberg, Christoph Dehio

**Author notes:** Address correspondence to Henning Stahlberg and Christoph Dehio. Authors contributed equally (Jaroslaw Sedzicki, Dongchun Ni).

## Abstract

The synthesis of complex sugars is a key aspect of microbial biology. Cyclic β-1,2-glucan (CβG) is a circular polysaccharide critical for host interactions of many bacteria, including major pathogens of humans (*Brucella*) and plants (*Agrobacterium*). CβG is produced by the cyclic glucan synthase (Cgs), a massive multi-domain membrane protein. So far, its structure as well as the mechanisms underlining the synthesis have not been clarified. Here we use cryo-electron microscopy (cryo-EM) and functional approaches to study Cgs from *A. tumefaciens*. We were able to determine the structure of this complex protein machinery and clarify key aspects of CβG synthesis. Our research opens new possibilities for combating pathogens that rely on polysaccharide virulence factors and can lead to new synthetic biology approaches for producing complex cyclic sugars.

## Introduction

Synthesis of different types of polysaccharides is critical for the biology of microorganisms. They constitute integral parts of the bacterial cell (e.g., peptidoglycan), are exposed on the bacterial surface (e.g., lipopolysaccharide, capsule components) or are secreted to the environment (e.g., glycans constituting the matrix of biofilms). Many polysaccharide virulence factors allow bacteria to invade their hosts and persist in them.

Cyclic glucans are produced by a number of Gram-negative bacteria^1^. Their structure, properties and function vary. In *Enterobacteria*, the cyclic enterobacterial common antigen plays a role in maintaining the outer membrane permeability barrier^2^. *Pseudomonas*, cyclic β-1,3-glucans were shown to sequester antimicrobial molecules within biofilms, leading to increased antibiotic tolerance^3,4^.

In *Rhizobiales*, cyclic β-1,2-glucans (CβG) are required for the proper host colonization. *Rhizobiales* include a number of pathogens and symbionts characterized by complex interactions with a broad range of hosts. Examples vary from human infection by zoonotic pathogens from the genus *Brucella*^5,6^, to plant root colonization by pathogenic (*Agrobacterium*) and symbiotic (*Rhizobium*) species^7^. CβGs have been implicated in osmotic adaptation, host environment recognition, immune modulation and lipid raft remodeling^8^.

The CβG synthesis pathway is well-conserved across *Rhizobiales*. The production of the main chain occurs in the cytoplasm and is orchestrated by the cyclic glucan synthase (Cgs), a large, multi-domain membrane protein that contains several enzymatic sites (Fig. 1a, Extended Data Fig. 1a). The mechanism driving CβG synthesis has only been partially deciphered, and many key aspects are not understood. The following steps have been proposed: (1) initiation involving the autoglycosylation of Cgs; (2) elongation of the glucan chain by the addition of Glc from UDP-Glc; (3) length control of the sugar chain performed by the C-terminal phosphorylase domain; and (4) cyclization that requires an intramolecular transglycosylation reaction^9–12^. The determination of Cgs structure and the interactions between different domains is critical for providing a comprehensive model. The mechanistic understanding of CβG synthesis can lead to new strategies of controlling microbes that rely on polysaccharide virulence factors. From the synthetic biology perspective, it may result in new approaches for generating complex sugars with desired properties.

**Fig. 1.**
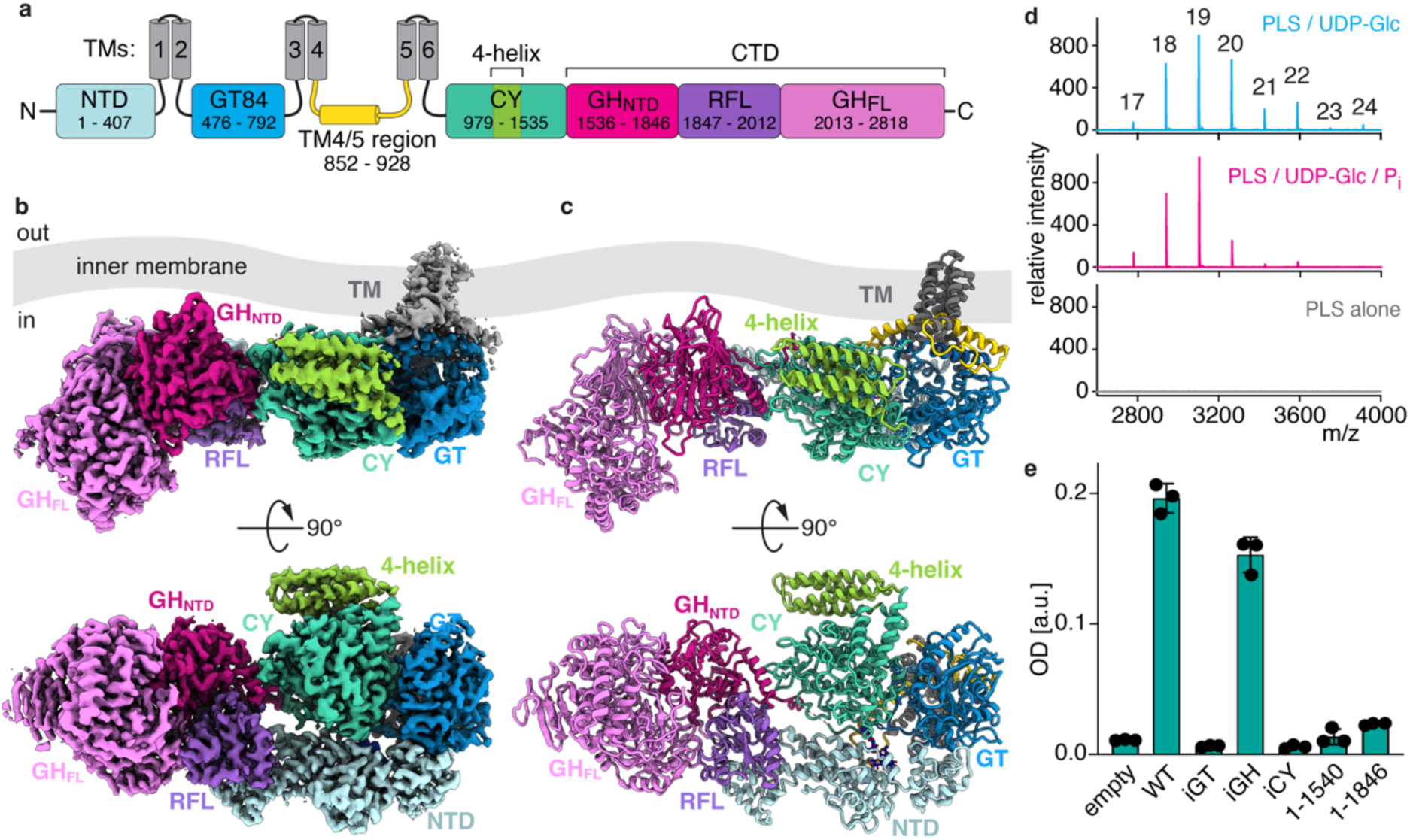
Overview of the Cgs structure and role of individual domains. **a,** Domain topology of Cgs. Cgs is embedded in the inner membrane with six TM helices (TMs 1-6). The soluble domains of the protein are all located on the cytoplasmic side of the membrane. They include the N-terminal domain (NTD, pale blue), a GT84 glycosyltransferase domain (blue), putative cyclase domain (teal) and two GH94 phosphorylase homology domains separated by a Rossmann fold-like domain (RFL, purple): GH94 N-terminal domain (GH94_NTD_, magenta) and full-length GH94 (GH94_FL_, pink). In addition, there is an extended stretch of 76 residues between TM helices 4 and 5 (yellow). The range of residues corresponding to different domains is indicated. **b-c,** Cryo-EM map (b) and model (c) of the full-length Cgs_iGH_. Side (top) and cytoplasmic (bottom) views are shown. Coloring scheme same as (A). The protein can be divided into two major parts: the NTD / GT / CY region located in the proximity of the transmembrane domain (TM, gray) and (2) a rigid CTD fold. The CTD contains three domains: partial GH domain (GH_NTD_, dark pink), a Rossmann fold-like fold domain (RFL, orange) and full-length GH domain (GH_FL_). The region between TMs 4 and 5 (yellow) forms a helix/loop motif at the TM-GT interface. **d,** MALDI-TOF analysis of CβG synthesis by Cgs proteoliposomes. The characteristic size distribution of glucan sizes can be observed. The number of Glc molecules within different cyclic glucan sizes is indicated. Addition of phosphate in the sample buffer (magenta) results in the synthesis of shorter glucan chains, indicating that the length control depends on the activity of the GH phosphorylase. **e,** Hypoosmotic growth assay showing the role of different domains of Cgs in functionality. The *Δcgs* mutant was complemented by plasmid-encoded *cgs* variants. Mutation of the active sites of GT and CY enzymatic domains (iGT, iCY) abolish activity. While the catalytic residues of the GH domain are not essential (iGH), truncation of the entire CTD (Cgs_1-1587_) or its part (Cgs_1-1846_) leads to loss of activity.

In this study, we present a series of cryo-electron microscopy (cryo-EM) structures of full-length Cgs from *A. tumefaciens* (*Atu*) in conformational states relevant to its physiological function. We were able to characterize the architecture of this complex, multi-domain glucan synthesis platform and provide insights into its enzymatic activities and regulation. Our results reveal a unique mechanism that uses a tyrosine-linked oligosaccharide as an intermediate in cycles of polymerization and processing of the glucan chain. Our work improves the understanding of complex polysaccharide synthesis and can potentially provide a basis for new approaches for fighting pathogens that rely on cyclic glucans as virulence factors.

## Results

### Cryo-EM analysis reveals the full-length structure of Cgs

According to previous studies^12–14^, CβG synthesis involves the cooperative action of two enzymatic domains performing chain elongations and length control. These are a GT84-family glycosyltransferase and a C-terminal GH94-family phosphorylase. The GT domain is believed to additionally initiate glucan synthesis by autoglycosylating Cgs at an unknown residue^12^. A putative third domain, referred to as the “cyclase” (CY), has been implied in the cyclization of the glucan chain, but its identity remains unknown (Fig. 1a).

In order to determine the structure of Cgs, we constructed a strain of *A. tumefaciens* C58 carrying an insertion in the chromosomal copy of the *chvB* gene encoding a C-terminal 3xFlag-tag (Cgs_WT_). In addition, catalytically inactive mutants of the GT (Cgs_iGT_) and GH (Cgs_iGH_) domains were generated by introducing substitutions of the predicted active site residues (D624A/D626A/D739A and D2393A/D2528A, respectively).

The resulting three constructs were purified, reconstituted into lipid nanodiscs^15^ and analyzed by cryo-EM in the presence or absence of UDP-glucose, leading to a number of distinct density maps (for an overview, see Extended Data Fig. 1b, 8-10). The Cgs_WT_ and Cgs_iGH_ samples produced high resolution maps of the full-length protein. The Cgs_iGH_ mutant displays reduced flexibility compared to WT, which resulted in improved cryo-EM maps obtained from much smaller datasets. All structures display a similar domain arrangement, but show some conformational changes. The most significant distinction between individual maps is the localization of densities corresponding to the UDP-Glc molecules as well as partial densities of glucan chain intermediates at different sites. These most likely represent different steps of the CβG synthesis cycle. In addition, a subpopulation of Cgs_iGH_ forms a homodimer, which allowed us to determine the Cgs_iGH3_ dimer structure. Finally, the Cgs_iGT_ mutations resulted in a partially disordered protein that only gave rise to a low-resolution map.

### Domain architecture of Cgs

Our data reveal a multi-domain structure embedded in the membrane through a small, 6-helix domain. The protein has an elongated shape that extends perpendicular to the TM domain, suggesting that in the native membrane-embedded state it is aligned along the lipid bilayer. The soluble part of the protein is divided into 4 distinct subdomains, all located on the cytoplasmic side (Fig. 1a-c).

The N-terminal domain (NTD) is characterized by an elongated fold that forms a side scaffold interacting with all the remaining parts of the protein. The GT84 domain is docked directly below the TM helices, resembling to GT2 family glycosyltransferases^13^. Residues 980-1539, corresponding to the putative cyclase domain (CY), form a globular domain that sits at the center of the structure. The C-terminal half of Cgs (CTD, starting with residue 1540) forms a rigid structure that can be divided into three distinct domains. The C-terminal end (residues 2020-2818) encodes for a full-length, enzymatically active GH94 family phosphorylase domain (GH_FL_) ^14^. Residues 1540-1846 form a domain corresponding to the N-terminal half of the GH94 domain fold (GH_NTD_). Both domains are separated by a small Rossmann fold-like domain (RFL, residues 1850-2012) that interacts with the NTD^16^.

### The CY domain is homologous to GH144 family endoglucanases

The fold of the CY domain indicates homology to a recently identified hydrolase family GH144 isolated from *T. funiculosus*^17^ (Fig. 1b, c, Extended Data Fig. 2a, b), which displays endoglucanase activity towards β-1,2-glucans. Based on the structure homology, we were able to identify the active site residues D1104/E1300 (Extended Data Fig. 2b). The Cgs_WT_ map contains a glucan density docked at the active site of CY in a manner similar to the soluble homolog. The homology to endoglucanases suggests that CY is capable of catalyzing a transglycosylation reaction of linear glucans, resulting in a cyclic chain. Compared to hydrolysis, the acceptor water molecule is replaced by a hydroxyl group belonging to the same glucan chain, leading to the cyclization and release of the cyclic sugar from the protein^18^.

The CY domain contains an additional stretch of around 120 residues absent in the soluble homolog. It forms a 4-helix motif protruding to the outside of the structure. The Cgs_iGH3_ model shows that this motif participates in the formation of a back-to-back dimer through an interaction with the CTD from a second Cgs molecule (Extended Data Fig. 2a, c). The dimer constitutes a small population of particles in the datasets. Interestingly, the TM domains of the dimerized Cgs molecules are at an angle of ∼90° relative to each other, indicating that the interaction could introduce curvature to the aligned inner membrane.

### Significance of the various Cgs domains in CβG synthesis

Previously reported Cgs activity assays were based on whole bacteria membrane fractions, and it has been unclear if the enzyme alone is sufficient for synthesizing CβGs. To exclude the role of other, unknown factors, we reconstituted purified Cgs into proteoliposomes (PLS) and performed an in vitro activity assay. MALDI-TOF analysis of soluble fractions confirmed that Cgs PLS can convert UDP-Glc into CβG (Fig. 1d, Extended Data Fig. 3). The presence of phosphate ions in the reaction buffer resulted in shorter glucan species, confirming that the length control depends on the phosphorylase activity of the GH domain ^11^. Surprisingly, Cgs PLS also display activity towards linear glucan chains. Addition of short linear β-1,2-glucan chains to the PLS resulted in the production of glucans with increased molecular weight (Extended Data Fig. 3c). This activity was independent of UDP-Glc and most likely indicates the transglycosylation activity proposed for the CY domain.

We wanted to further test the importance of the three enzymatic domains of Cgs for the function. To this end, we used the hypoosmotic growth assay, which is based on the sensitivity of CβG-deficient *Atu* to low osmotic stress (Fig. 1e, Extended Data Fig. 4) ^19^. A *τ1chvB* mutant of *Atu* was generated and different constructs of the *chvB* gene were tested for their ability to rescue the deletion phenotype. The results clearly indicate that enzymatically active GT and CY domains are required for CβG synthesis. Inactivation of the GH enzymatic activity by replacement of two essential amino acids in the catalytic center (D2393A/D2528A) had only a minor impact on bacterial survival, suggesting that glucan length control is not critical for Cgs functionality, which is in accordance with previous studies^11^. The presence of the CTD domain itself, however, seems important, and its complete (Cgs_1-1587_) or partial truncation (Cgs_1-1846_) has a detrimental effect. This hints to a scaffolding role of the CTD that is independent of length-control.

### The GT84 domain displays a novel fold with partial GT2 family characteristics

The GT84 domain responsible for the synthesis of the linear glucan chain is located directly below the transmembrane domain (Fig. 2a, b, Extended Data Fig. 5). The structure of the domain and the arrangement relative to the TM helices resembles membrane-embedded family-2 GTs as exemplified by the cellulose synthase BcsA (Extended Data Fig. 5a) ^13^. Characteristic conserved motifs can be identified, including the D527/D624 that coordinate the UDP as well as the catalytic base D739 (BcsA D179/D246 and D343, respectively). There are, however, major differences. In Cgs, the TM domain is relatively compact and does not form an export channel. While the interface helices (IFs) 2 and 3 and the gating loop are present, the region homologous to IF1 (IF1h, residues 684-701) forms a loop that extends into a chamber at the interface between the NTD and CY domains. This rearrangement opens a channel from the GT active site that continues underneath the TM domain towards the NTD (Fig. 2a, Extended Data Fig. 5b). The channel does not contain a density corresponding to the linear glucan chain in any of the maps. The PilZ domain, responsible for binding of the second messenger cyclic-di-GMP and regulation of the gating loop in BcsA, is absent in Cgs (Extended Data Fig. 5a).

**Fig. 2.**
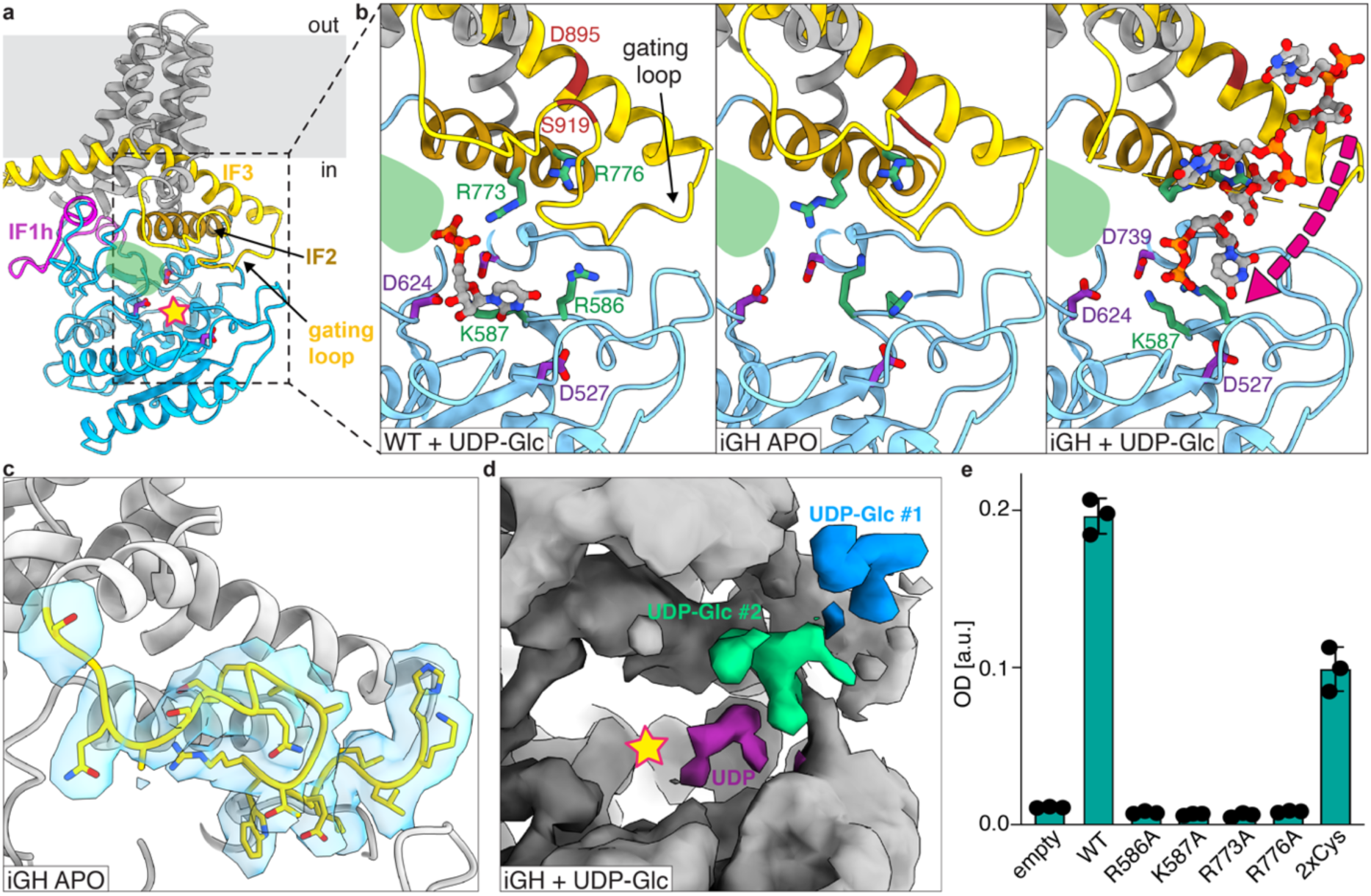
The glycosyltransferase domain. **a,** The GT domain (blue) is located below a compact TM domain (gray). IF1-homology (IF1h, magenta), IF2 (brown) and IF3 (yellow) motifs are located at the interface. The modification of the IF1 motif allows access to the other side of the domain through a channel underneath of the TM bundle (green shading). The GT active site is marked with a star. The NTD and CY domains are not shown for clarity. **b,** Close-up of the GT active site of three models: Cgs_WT_ (left) and Cgs_iGH1_ (middle) Cgs_iGH2_ (right). The Cgs_WT_ model contains a UDP molecule density docked at the active site, indicating a post-transfer state. In the presence of the substrate, the gating loop of Cgs_iGH_ is disordered and two UDP-Glc and one UDP (gray-orange sticks) are found at the IF2/IF3 motifs and near the active site. Residues R586, K587, R773 and R776 that facilitate substrate recruitment into the active site are indicated in green. Key active site residues are shown in purple. Residues D895 and S919 on IF3 that were substituted with cysteines are indicated in brown. The magenta arrow indicates the loading of the UPD-Glc molecules into the active site in the proposed model. **c,** Close up of the resting-state gating loop (yellow) in the Cgsi_GH1_ model and the corresponding density map (blue). **d,** Close up of the GT density in the Cgs_iGH2_ map. The densities corresponding to the two UDP-Glc molecules (blue, green) are bound to IF3. The UDP (purple) was found near the GT active site. **e,** Hypoosmotic growth assay showing the role of the positively-charged residues at the UDP-Glc-binding site. The *Δcgs* mutant was complemented by plasmid-encoded *cgs* variants. The double-cysteine mutant showed reduced activity compared to WT.

The Cgs_iGH_ models indicate that the presence of UDP-Glc affects the gating loop conformation (Fig. 2b-d). In the apo state, the loop can be found above the GT84 active site. In the presence of the substrate, the motif seems disordered, and there are additional densities bound in the proximity of the active site and the IF3. We assigned these densities as UDP-Glc and identified a potential binding site composed of positively-charged side chains R586/K587/R773/R776. The hypoosmotic stress assay indicates that single alanine substitutions of these residues were sufficient to completely abolish growth, indicating a key role (Fig. 2e). In addition, stabilizing the association between the gating loop and IF3 by introducing Cys residues (D895C/S919C) resulted in reduced function. This suggests that the substrate-induced displacement of the gating loop is required for synthesis to take place. We hypothesize a regulatory mechanism, in which an additional UDP-Glc binding site serves a dual function – it allows allosteric induction of the enzymatic activity by the substrate and at the same time recruits UDP-Glc to the proximity of the active pocket.

### Cgs autoglycosylates the IF1h loop to form the glucan primer

It was proposed that a short, covalently-linked glucan chain remains permanently associated to the protein and serves as a primer for elongation during each cycle of glucan synthesis. Although the GT84 domain was proposed to autoglycosylate Cgs (initiation step), the mechanism as well as the site of this modification have been unclear^12^. Our cryo-EM maps contain a polysaccharide density attached to the residue Y694, which is a part of the IF1-homology loop (Fig. 3, Extended Data Fig. 6). The density corresponds to a chain of minimum 8-9 glucose molecules that interacts with the tetratricopeptide (TPR) homology motif of the NTD. The IF1h loop and the O-glucan are located in a large chamber formed at the interface between the NTD, GT and CY domains, which is accessible from the cytoplasm through a large cavity (Extended Data Fig. 6c). The glycosylation of Cgs is essential and the Y694A mutation completely abolishes activity (Fig. 3b).

**Fig. 3.**
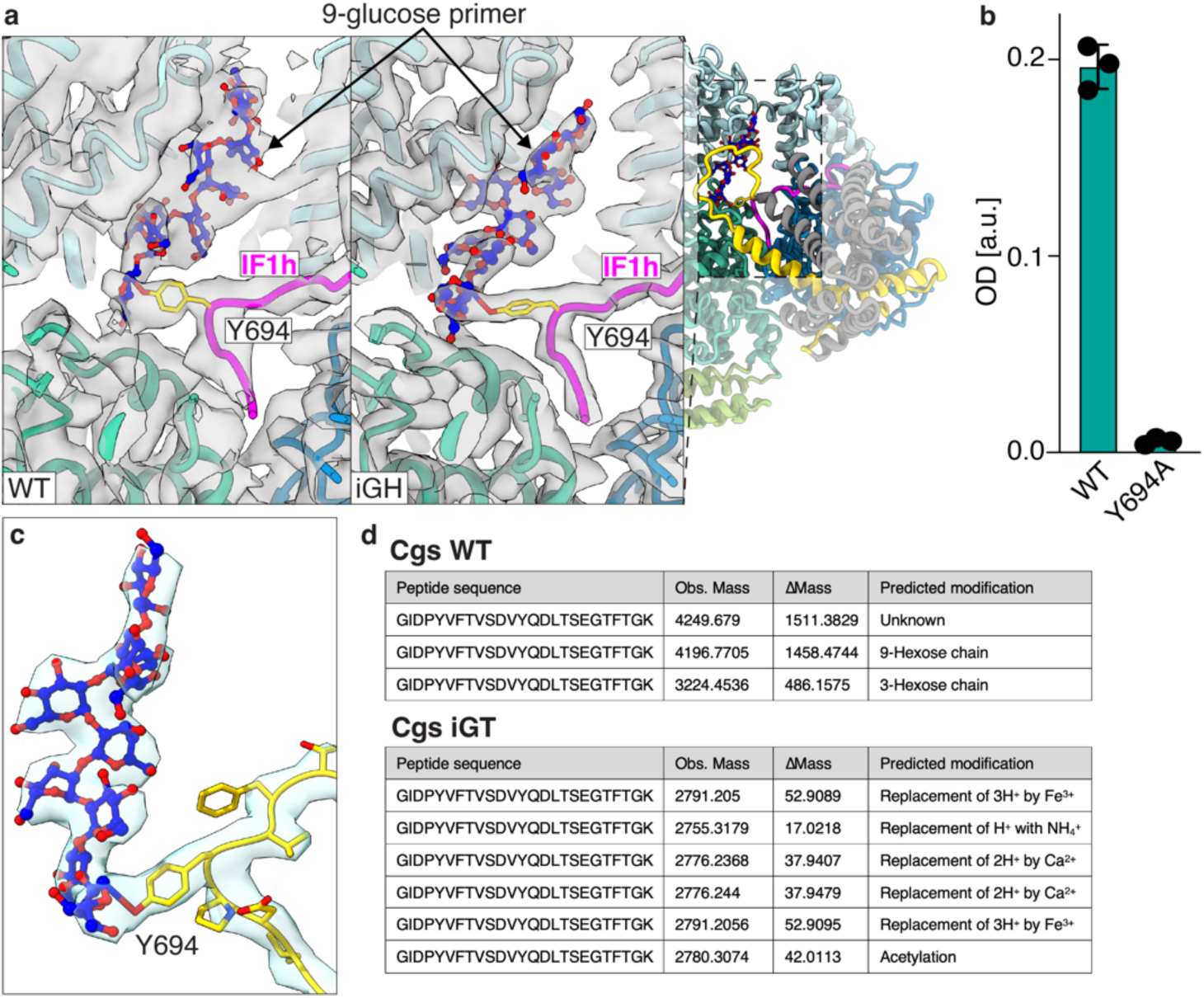
Cgs is O-glycosylated at the Y694 residue of the IF1h loop. **a,** Cgs seen from the side of the membrane. The glucan primer chamber formed between the NTD (light blue), GT (blue) and CY (teal) domains in Cgs_WT_ (left) and Cgs_iGH1_ (right) maps. The Y694 residue at the tip of the IF1-homology loop (magenta) is inserted into the chamber. The additional density originating form Y694 indicates a linear O-glycan chain (blue-red sticks) attached to the protein. The feature is best resolved in the maps obtained for the iGH mutant. **b,** Hypoosmotic growth assay indicated the essential role of Y694 in Cgs activity. **c,** Close-up of the IF1h loop (yellow) and glucan primer (blue-red) with corresponding density map (blue). **d,** Mass spectroscopy results of purified Cgs_WT_ and Cgs_iGT_. Peptides containing the Y694 glycosylation site were detected. The table list the observed masses (Obs. Mass) as well as the difference between the observed mass and the theoretical mass of the peptide (ΛMass). Modifications corresponding to 9- and 3-hexose chains were detected for Cgs_WT_, but were absent in Cgs_iGT_.

While O-glycosylation of Ser and Thr residues is well studied, there is sparse information regarding Tyr^20,21^. One example is glycogenin, which is capable of both catalyzing Tyr O-glycosylation as well as extension of a short glucan chain^20^. To confirm that the GT domain is O-glycosylating Y694, we used mass spectroscopy to assess the modification state of the residue in purified Cgs_WT_ and Cgs_iGT_ (Fig. 3d). We found that an additional mass corresponding to 9- and 3-glucose chains is attached to the IF1-homology loop in Cgs_WT_, while no such modifications were detected in Cgs_iGT_.

While our results clearly indicate the autoglycosylation of Cgs, Y694 is located relatively far from the GT active site. The low-resolution Cgs_iGT_ map indicates that in the absence of the O-glucan chain, the interface between the NTD and GT is disrupted, resulting in increased flexibility (fig. S6D). It is possible that this allows the access of the IF1h loop to the active site in newly-synthesized Cgs molecules, enabling glucan primer synthesis that leads to the mature conformation of the protein.

### The CTD forms a glucan binding landscape

The C-terminal half of Cgs constitutes a large, rigid fold (Fig. 4, Extended Data Fig. 7a, b). Two of its subdomains bear close resemblance to the GH94 family cellobiose phosphorylases^14^: a partial GH_NTD_, and an enzymatically active GH_FL_. This results in a pseudodimer interface that reflects the dimerization of the soluble homolog (Extended Data Fig. 7b). Both domains are separated by an additional small Rossmann fold-like domain (RFL), with a 4-strand β-sheet sandwiched between a number of α-helices^16^. Although the GH active site is not essential for the synthesis of CβGs, truncations of the CTD have a strong effect (Fig. 1d), indicating additional functions of the domain.

**Fig. 4.**
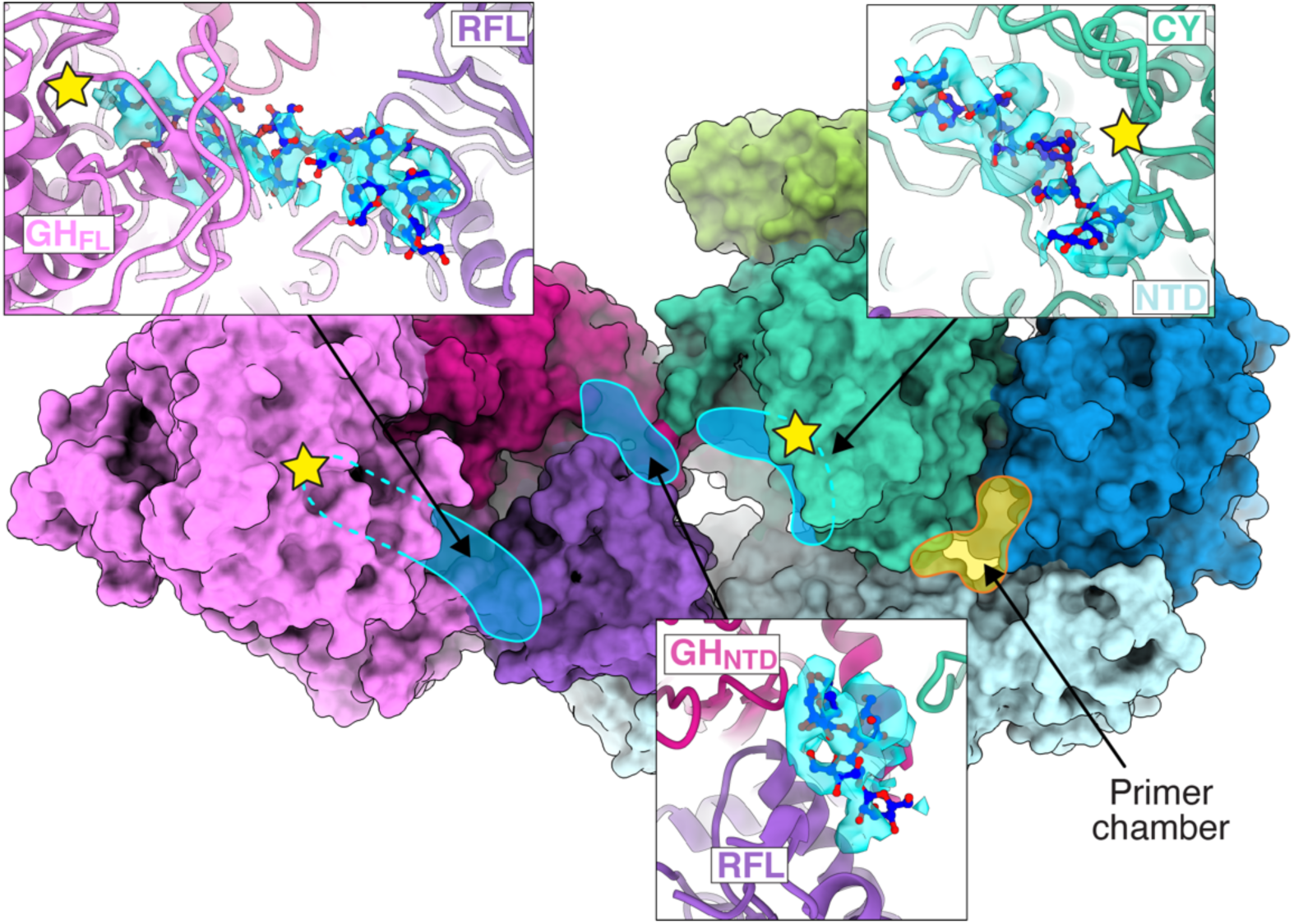
The CTD forms glucan-binding landscape for glucan processing. Cytoplasmic view of Cgs. The glucan-binding landscape constitutes a large surface between the active sites (yellow stars) of the CY (teal) and GH_FL_ (pink) that contains multiple glucan affinity sites (blue shading). The landscape allows the coordination of the glucan chain during the processing by the CY and GH domains. Side panels indicate glucan densities that were observed interacting with the surface in different maps: at the CY active site (Cgs_WT_ map, top right), at the GH_FL_/RFL interface, docked to the GH_FL_ active site (Cgs_iGH_ map, top left) and a less ordered density at the GH_NTD_/RFL contact site (Cgs_iGH_ map, bottom). Glucan chain fragments are shown as blue-red sticks.

The Cgs_WT_ and Cgs_iGH2_ maps obtained in the presence of UDP-Glc contain additional densities of glucan chain fragments bound to the protein at the cytoplasmic side of the CTD (Fig. 4). This surface, which stretches between the CY and GH_FL_ enzymatic pockets, appears to contain several affinity sites capable of coordinating the polysaccharide. They include a glucan chain docked to the CY active site (Cgs_WT_), and two binding sites that localize at the interfaces between RFL and neighboring domains (Cgs_iGH2_): GH_NTD_/RFL and GH_FL_/RFL. The glucan bound at the GH_FL_/RFL site ends within the active site of the GH domain. The second density, at the GH_NTD_/RFL interface, is shorter and less resolved. It is bound to the GH_NTD_ in a manner resembling the substrate interaction in the dimer of the soluble homolog (Extended Data Fig. 7b). We propose that this glucan binding landscape could provide a platform for ordered chain displacement, orienting the nascent polysaccharide for further processing that leads to cyclization by the CY domain.

### The N-terminal half of Cgs is highly dynamic

While the C-terminal half of Cgs is characterized by a rigid fold, the N-terminal half is highly dynamic (Extended Data Fig. 7c). In fact, most of the initially extracted particle datasets produced density maps with a well-defined CTD and weak partial density corresponding to the rest of the protein. The overall flexibility is reduced to some extent in the iGH construct, suggesting that at least some of the movement is related to the processing of the glucan chain at the active site. Our full-length maps likely represent relatively low energy states of the protein.

The 3D variability analysis of the full-length structures indicates different kinds of rearrangements. Most prominent ones include a hinge movement at the N- and C-terminal halves of the structure, as well as changing distance between the NTD and CY domains. In the latter case, the movement produces an opening between the primer chamber and the glucan binding landscape. This opening likely allows the passage of the sugar chain at certain steps of the synthesis cycle.

## Discussion

Cgs constitutes a self-contained, dynamic assembly line. The synthesis of CβGs is a multi-stage process that requires the coordination of several enzymatic activities in time and space. Our work reveals the key parts of this machinery, including the enzymatic domains as well as multiple motifs that allow the proper positioning and migration of the glucan chain. Our data indicate that Cgs is highly dynamic. The movement of the different parts of the protein relative to each other is apparent, especially between two halves of the molecule. This high flexibility could play a role during CβG synthesis, and there possibly are conformations important for the function that are dynamic and difficult to capture by cryo-EM.

Our work identifies the Y694 glycosylation site and provides evidence supporting the dual role of the GT domain, which facilitates both the extension of the glucan as well as the autoglycosylation that creates the glucan primer. In fully folded Cgs, the access of Y694 to the active site is problematic. At the same time, the lack of modification in the iGT mutant seems to destabilize the entire protein. In newly produced Cgs polypeptides, this increased flexibility could facilitate the addition of the O-glucan primer to Y694, leading to a synthesis-ready conformation. The autoglycosylation likely occurs only once during the protein’s lifetime and the glucan primer is reused in subsequent cycles.

The requirement for a covalent link between the glucan chain and Cgs distinguishes it from previously described membrane-embedded GTs^13,22,23^. The glucan primer can facilitate synthesis initiation and stabilize protein-sugar interactions in further steps, preventing dissociation during conformational rearrangements. While O-glycosylation of Ser and Thr residues is well studied, only sparse examples of enzymes capable of Tyr O-glycosylation are known. So far, only glycogenin was shown to combine this function with sugar chain extension^20^, and Cgs is the first example of this bifunctionality in bacteria.

The substrate-induced conformational changes of the GT gating loop hint to an alternative regulatory mechanism employed in the absence of the PilZ-domain, which could be explored as a potential drug target. Molecules stabilizing the gating loop in the inactive state could potentially have a strong inhibitory effect on Cgs, and with that act as potent antimicrobials against pathogens allowing that rely on cyclic glucans. These findings can be potentially extended to related membrane-bound family 2 GTs^22,23^.

The close structural resemblance of the CY domain to soluble endoglucanases from the GH144 family indicates that cyclization involves the intramolecular transglycosylation of the GT domain product. In this reaction, enzymes displaying a hydrolase fold cleave a glucan and use a hydroxyl group from a different part of the same chain as an acceptor. This process requires precise coordination of the glucan relative to the active site^18^. In Cgs, the binding landscape formed by the subdomains of the CTD provides an affinity platform that arranges the chain in an appropriate conformation, allowing length control and subsequent cyclization. Additionally, the CTD is responsible for the dimerization through interacting with the 4-helix motif of the CY. However, the exact role of the dimer remains unclear. The interaction could stabilize certain conformations of individual Cgs monomers or provide cues between different domains to control their sequential action.

Based on our findings, we propose a general model for CβG synthesis (Fig. 5), which can be separated into 2 major stages: synthesis (driven by GT) and processing (driven by CY and GH). The 9-glucose O-glucan primer attached to the IF1h loop is synthesized by the GT during Cgs folding and is reused in subsequent cycles. In the stable resting state, a primer is interacting with the TPR-homology part of the NTD (step 1). A large cavity between the NTD, GT and CY domains allows access from the glycosylation site to the surrounding solvent. The synthesis requires the transition of the non-reducing end of the primer into the GT active site (step 2). Based on structures of related GTs, this most likely occurs through the channel created by the unwinding of the IF1h. We were unable to observe the glucan chain in the channel in all obtained structures. This could be caused by a number of reasons, including the flexibility of the protein and instability of this conformational state. It is also possible that the affinity of the channel for the glucan chain is relatively low, leading to detachment during sample preparation. The elongation of the sugar chain requires the interaction of UDP-Glc with the IF3, which displaces the gating loop and allows the positioning of the substrate in the proximity of the enzymatic pocket.

**Fig. 5.**
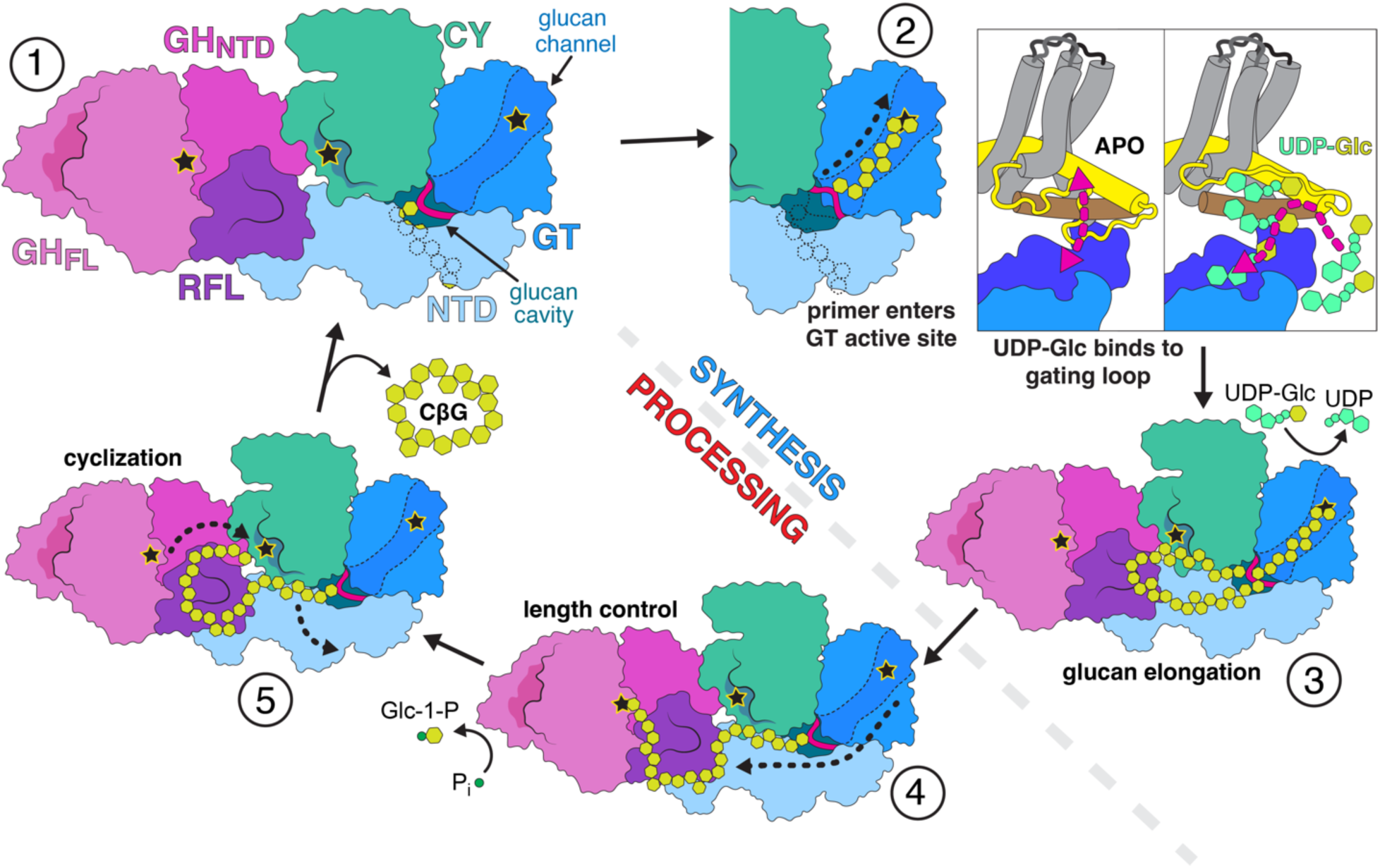
Proposed mechanism of cyclic glucan synthesis by Cgs. The Cgs cycle can be separated into two major stages: glucan synthesis (driven by the GT domain) and processing (driven by both CY and GH). Coloring of individual domains is indicated. Active sites of enzymatic domains are marked with stars. In the resting state (1), the primer resides in the primer chamber and is interacting with the NTD. The synthesis requires a number of rearrangements of the glucan chain (black dashed arrows). (2) The synthesis requires the entry of the non-reducing end of the primer into the GT active site. Glucan chain synthesis is regulated by the concentration of the UDP-Glc substrate, which affects the conformation of the gating loop. The elongation of the chain (3) continues until its length allows contact with the CTD surface, which leads to the translocation of the non-educing end of the glucan and length control. (5) The glucan affinity sites allow orient the glucan chain in a conformation that promotes an intrachain transglycosylation, leading to the release of a cyclic glucan molecule. The reaction regenerates the primer, which can be re-used in a subsequent cycle. Dashed gray line separates the synthesis and processing stages of the cycle. See main text for a detailed description.

The glucan primer is extended by the addition of new glucose molecules to the non-reducing end (step 3). The growing sugar chain forms a loop that is pushed through the cavity. Once the chain reaches a certain critical length, it interacts with the CTD, triggering glucan displacement and the transition to the processing stage. Although the exact mechanism behind the switch is difficult to deduce, it results in the coordination of the glucan chain at the binding landscape, with the non-reducing end accessible to the GH active site (step 4). The length control can occur at this point, but it does not seem essential for cycle completion. Finally, CY catalyzes an intramolecular transglycosylation, which releases a cyclic CβG molecule and restores the 9-glucose primer (step 5).

Future work is required to fully clarify the exact mechanism leading to the synthesis of a CβG chain. Insights into the process of CβG synthesis can provide new tools to combat pathogens that use polysaccharides as virulence factors. It can also equip us with new approaches for the synthesis of complex polysaccharides using biotechnology approaches.

## Materials and Methods

### Plasmids

For a detailed description of all plasmids used in this study, see Table S3. DNA ligations were performed using the In-Fusion HD Cloning Kit (Takara Bio). The identity of all constructs and mutant strains was confirmed by sequencing (Microsynth AG). See Table S2 for the complete list of primers and their specific purpose. The derivatives of pNPTS138 used for directed mutagenesis were generated by amplifying chromosomal DNA fragments from heat-killed *Atu* colonies and ligating them into the pNPTS138 backbone digested with SalI and HindIII. The pJS801 used in the hypoosmotic growth assay is a derivative of pBBR1 generated by amplifying the *chvB* gene sequence with the upstream promoter region from heat-killed *Atu* using primers pJS1108/prJS713. The PCR product was ligated with the pBBR1MCS-2 amplified with primers prJS751/prJS1072. The derivatives of pJS801 carrying point mutations were generated by amplifying the *chvB* gene sequence using primers introducing relevant point mutations (see Tables S1 and S2) and re-ligating them with pBBR1.

### Bacteria strains

For a detailed description of all strains used in this study, see Table S4. Stellar competent cells (Takara Bio) provided with the In-Fusion Cloning Kit were used in all cloning procedures. All strains were maintained in lysogeny broth (LB, 10 g l^-1^ tryptone, 5 g l^-1^ yeast extract, 10 g l^-1^ NaCl). When required, kanamycin (50 μg ml^-1^), oxytetracycline (12 μg ml^-1^ for *E. coli* or 3 μg ml^-1^ for *Atu*) and 5-aminolevulinic acid (50 μg ml^-1^) were added to growth media. All *E. coli* strains were grown at 37°C unless stated otherwise. *A. tumefaciens* C58 strains were cultivated at 28°C. Mutagenesis vectors were introduced into *Atu* strains by conjugation using the ST18 donor strain ^24^. Expression plasmids were introduced by electroporation. All chromosomal modifications of the *Atu* C58 were achieved with derivatives of the pNPTD138 suicide vector. Following conjugation into the target strain, clones that have undergone two successful recombination events were selected in a two-step process. The required genomic changes in the targeted loci were confirmed by sequencing. The *Atu* strain used in the growth assay is a derivative of *Atu* C58 with a deletion of the *cgs* gene (*ΔchvB, Atu004*) using plasmid pJS301. Rescue plasmids carrying different constructs of *cgs* were introduced into *Atu004* to generate strains used in the hypoosmotic growth assay. The strain used for expressing the Cgs-Flag construct was generated by introducing a genomic 3xFlag-tag sequence to the end of the *cgs* (*chvB*) gene using plasmid pJS305. The resulting *Atu009* strain was later used to generate two additional expression strains carrying mutations in the active sites of GT84 (*Atu130*, generated using plasmid pJS091) and GH94 (*Atu125*, generated using plasmid pJS309). The expression of the *cgs-3xFlag* constructs in the resulting *Atu* strains was confirmed by Western blot using an anti-Flag antibody (Sigma).

### *Atu* growth assay

*Atu* strains were precultured overnight at 28°C in 14 ml polypropylene tubes (Sarstedt) in a rotary shaker. The OD_600_ of the cultures was measured in a BioPhotometer #6131 (Eppendorf). The cultures were then used to inoculate YPL medium (0.1% peptone, 0.1% yeast extract, 0.09% glucose) at a final OD_600_ of 0.01. The cultures were then dispensed into 96-well flat bottom plate (160 μl per well) and placed Synergy H4 Hybrid Microplate Reader (BioTec) and incubated with fast shaking at 28°C for indicated periods of time. OD_600_ was measured every 60 min. Technical well duplicates were used for each strain. Each reported result represents at least 3 (n=3) biological replicates.

### Cgs expression and purification

*Atu* strains carrying the chromosomal 3xFlag insertion at the C-terminus of the *chvB* gene (Atu009, Atu125, Atu130) were grown at 28°C in LB medium until OD_600_ of 4.5-5.5. The cells were collected by centrifugation and frozen at −80°C until further processing. All purification steps were performed at 4°C. Bacteria pellets were resuspended in lysis buffer (50 mM HEPES pH 7.5, 500 mM NaCl) supplemented with cOmplete protease inhibitor tablets (Roche) and lysed by 3 passes at 20’000 psi through a microfluidizer (Microfluidics). Large debris was removed by centrifugation (30 min, 10,000g) and cell membranes were pelleted by ultracentrifugation (45 min, 200’000g). Membrane pellets were solubilized with 1% n-dodecyl-β-d-maltopyranoside (DDM, Anatrace), loaded onto Flag affinity agarose beads (Sigma) and washed extensively with buffer containing 0.05% DDM. Following elution with 0.4 mg/mL 3xFlag peptide (Sigma), the samples were purified by size-exclusion chromatography on a Superose 6 Increase 10/300 column (GE Healthcare).

### Reconstitution of Cgs into nanodiscs

*E. coli* polar lipid extract (Avanti Polar Lipids) was solubilized in chloroform, dried under nitrogen gas to form a lipid film, and stored under vacuum overnight. The lipid film was resuspended at a concentration of 25 mM in buffer containing 20 mM HEPES, pH 7.5, 150 mM NaCl and 300 mM sodium cholate. Purified Cgs, the MSP1D1 membrane scaffold protein^15^ and lipids were mixed at a molar ratio of 1:4:100 in buffer containing 25 mM HEPES, pH 7.5, 150 mM NaCl and incubated for 30 min. at 4°C. Detergents were removed by incubation with 100 mg Bio-Beads SM2 (Bio-Rad) overnight at 4°C. The Cgs-nanodisc complexes were purified using a Superose 6 Increase 10/300 column (GE Healthcare) in a buffer containing 25 mM HEPES, pH 7.5, and 150 mM NaCl.

### Reconstitution of Cgs into proteoliposomes

*E. coli* polar lipids at a concentration of 10 mg ml^-1^ in 25 mM HEPES pH 7.5, 100 mM NaCl were extruded through a 100 nm filter to generate liposomes. Purified Cgs at a concentration of 2 mg ml^-1^ was added to liposomes destabilized by 0.3% DDM at a 1:2’000 protein:lipid molar ration. Detergent was then removed by two rounds of fresh Bio-Beads. Resulting proteoliposomes were loaded onto a Sephadex G50 column equilibrated with buffer 1 (25 mM Tris pH 7.6, 100 mM NaCl). Concentration of protein in the eluted fractions was measured using the Bradford method (protein assay dye purchased form Bio-Rad). Fractions containing the proteoliposomes were combined and adjusted to 0.2 mg ml^-1^ protein concentration. The incorporation of Cgs into proteoliposomes was confirmed by SDS-PAGE.

### CβG in vitro synthesis assay

Cgs proteoliposomes were diluted to 0.1 mg ml^-1^ protein concentration in buffer 1. The final concentrations of reaction components were 0.1 mg ml^-1^ protein, 5 mM UDP-Glc, 5 mM MgCl_2_, 5 mM sodium phosphate, 5 mg ml^-1^ linear β-1,2-glucan (Megazyme) in 300 μl final sample volume. After pipetting on ice, reactions were incubated for 24 h at 22°C with slow rotation. Afterwards, the liposomes were removed by running the sample through 100 kDa cutoff Amicon centrifugal filters (Merck). Finally, the sample buffer was exchanged to water in 3 kDa cutoff Amicon centrifugal filters (Merck). The samples were then concentrated to a final volume of 30 μl and subjected to mass analysis. Mass spec analysis of glucans

### MALDI-TOF mass spectroscopy analysis

Samples were measured under conditions optimized for low mass materials (0.5-6 KDa, for the detection of glucans). 2,5-dihydroxybenzoic acid (DHB, Sigma-Aldrich) was used as the matrix compound and 50 mg/mL DHB was dissolved in a 1:500:500 v/v/v TFA/water/acetonitrile solution and then mixed with the sample in a 1:1 volume ratio. The samples were mixed. 1.5 μL of each sample was spotted on a stainless steel MALDI plate and dried in air. Mass spectrometric data were collected in reflectance mode (RP) ion mode on a MALDI-TOF/TOF mass spectrometer (Bruker AutoFlex Speed). MALDI was launched by a Nd:YAG SmartbeamTM-II 2 kHtz laser (355 nm) with 84% of intensity per spectrum, accumulated 1000 shots.

### Electron microscopy sample preparation

Purified, nanodisc-reconstituted protein samples were concentrated to 2.0-2.5 mg ml^-1^. To generate substrate-bound samples, the protein was incubated with 5 mM UDP-Glc and 1.5 mM MgCl_2_ at 22°C for 2 h directly before freezing. 4 μl of protein solution was applied to glow-discharged Quantifoil holey carbon grids (2/2, 300 mesh). Grids were blotted for 3 s at 100% humidity and 4°C, using a Vitrobot Mark IV (ThermoFischer).

### Cryo-electron microscopy data acquisition and processing

For the Cgs_WT_ sample, cryogenic grids were screened on a Glacios TEM (TFS 200 kV) to determine the ice quality and then transferred to a Titan Krios G4 TEM (TFS 300 kV) equipped with Cold-FEG and a Falcon IV detector. EPU v2.12.1 (TFS) software was used for data automation. The range of defocus at exposure was −0.8 to −2.5 µm and the total dose calibrated was 50 e/Å2. Data were exported to EER format. For the Cgs_iGH_ sample, movies were collected on a Titan Krios (TFS) operating at 300 kV, equipped with a Gatan Quantum-LS energy filter (20 eV zero-loss energy filter) and a K2 Summit direct electron detector (Gatan Corporation, California, USA). SerialEM was used for data automation^25^. Dose fractionated images were acquired in counting mode with a nominal magnification of 165 kx, corresponding to a physical pixel size of 0.82 Å. The defocus range at exposure was 1.0 to −2.2 µm. Detailed data acquisition statistics and image processing workflow are shown in figs. S9-11 and Table S1. Image processing was performed with cryoSPARC v3.4^26^. Raw movies were motion corrected, dose weighted, pre-sorted and exported in mrc format. For the Cgs_WT_ sample, 21,835 movies were imported into cryoSPARC v3.4. The contrast transfer function (CTF) was estimated using the cryoSPARC patch CTF implementation as well as Ctffind^26,27^. To create a 2D template, blob picking was applied to 3000 micrographs and then 2D classification was performed. Using the template picking implementation from cryoSPARC, 2,181,584 particles were then automatically populated. Due to the obviously dominant orientations of the particles, 2D rebalancing was crucial for the 2D classification process, especially with manual reduction of particles with top and bottom views. After six rounds of 2D classifications, 225,741 particles were selected and Ab-Initio & Hetero Refinement was performed, yielding an optimal 3D class with 149,348 particles. Moreover, using cryoSPARC 3D classification, the particle subset was divided into seven classes. By combining class 4 and class 5, 49 830 particles were implemented with 3D refinement and a 3D map with a global resolution of 3.48 Å was obtained. To improve the quality of the glucan binding to the CY domain, subsequent local refinement was performed with a conventional soft mask volume surrounding the CY domain area. The final cryoEM map resolution was measured via gold standard method FSC with a cutoff value of 0.143. The local resolution distribution was estimated by the Local. Res. implementation in cryoSPARC. For the Cgs_iGH_ samples, the data were processed in cryoSPARC v3.4 in a manner similar to WT samples. Data processing started with particle picking, 2D classification, ab-Initio along as well as Hetero Refinement and 3D classifications have been applied, the best 3D classes were subjected to 3D refinement and local refinement. However, a slight difference is that the flexibility and dominant orientation issues are significantly better for the Cgs_iGH_ samples compared to Cgs_WT_. In addition, the 2D classes of Cgs dimers could be clearly identified from the 2D classification and the dimers could be reconstructed in 3D, as shown in fig. S9.

### Model building and refinement

Resolutions and density qualities of the obtained cryoEM maps were sufficient for de novo model building. The initial model was built in coot v0.9.4.1 by manually tracing the backbone^28^. The sequence was then assigned and matched the densities. After several rounds of manually building and adjusting the model, the geometry, model clashes, and rotamers were optimized in Phenix v1.19.2-4158 with real_space_refine plugin^29^. The relevant statistics are in Table S1. Figures were prepared with UCSF Chimera and UCSF ChimeraX v 1.4^30,31^.

#### Proteomics sample preparation

Purified Cgs_WT_ and Cgs_iGT_ protein samples were heated to 95° C for 10 min. Proteins were alkylated using 15 mM iodoacetamide at 25° C in the dark for 30 min. For each sample, 50 µg of protein lysate was captured, digested (trypsin 1/50, w/w; Promega), and desalted using STRAP cartridges (Protifi) following the manufacturer’s instructions.

### LC-MS analysis of immunoprecipitated samples

Approximately 250 ng of peptides were subjected to LC–MS/MS analysis using a Q Exactive HF Mass Spectrometer fitted with an Ultimate 3000 (both Thermo Fisher Scientific) and a custom-made column heater set to 60°C. Peptides were resolved using a RP-HPLC column (75μm × 30cm) packed in-house with C18 resin (ReproSil-Pur C18–AQ, 1.9 μm resin; Dr. Maisch GmbH) at a flow rate of 0.3 μL min^-1^. A linear gradient of buffer B (80% acetonitrile, 0.1% formic acid in water) ranging from 7% to 30% over 30 minutes and from 30% to 50% over 5 minutes was used for peptide separation. Buffer A was 0.1% formic acid in water. The mass spectrometer was operated in DDA mode with a Top5 method. Each MS1 scan was followed by high-collision-dissociation (HCD) of the 5 most abundant precursor ions with dynamic exclusion set to 20 seconds. For MS1, 3e6 ions were accumulated over a maximum time of 100 ms and recorded at a resolution of 120’000 FWHM (at 200 m/z). MS2 scans were acquired at a target setting of 1e6 ions, maximum accumulation time of 200 ms and a resolution of 60’000 FWHM (at 200 m/z). Singly charged ions and ions with unassigned charge state were excluded from triggering MS2 events. The stepped normalized collision energy was set to 25, 30 and 35. The mass isolation window was set to 1.4 m/z and one microscan was acquired for each spectrum.

### Proteomics data analysis

The raw files were converted to mzML format by MSConvert in conjunction with ProteoWizard^32^. The mzML files were analyzed using FragPIpe (18.0) ^33^ with settings for open search^34^. In brief, the spectra were searched against a database containing the protein of interest and common contaminants and their decoys using the following search criteria: full tryptic specificity was required (cleavage after lysine or arginine residues, unless followed by proline); 2 missed cleavages were allowed; carbamidomethylation (C) was set as fixed modification; oxidation (M), phosphorylation (STY) and acetylation (Protein N-term) were applied as variable modifications; mass tolerance of −2 m/z to 2000 m/z on precursor level.

## Acknowledgments

We thank the BioEM Lab (University of Basel, Switzerland) and the Dubochet Center for Imaging (EPFL, University of Lausanne and University of Geneva) for the cryo-EM data collection. We thank C. Perez and G. Cebrero Acuña (University of Basel, Switzerland) for their help with proteoliposome generation and handling. We thank Daniel Ortiz (from mass spectrometry and elemental analysis platform in EPFL) for the technical assistance provided for mass spectrometry analysis. We also thank L. Siewert for help during manuscript preparation.

## Data Availability

Atomic structure coordinates were deposited in the Protein Data Bank under accession codes PDB:XXXX (Cgs_WT_), PDB:XXXX (Cgs_iGH1_), PDB:XXXX (Cgs_iGH2_), PDB:XXXX (Cgs_iGH3_). The cryo-EM maps were deposited in the Electron Microscopy Data Bank under accession codes EMD:XXXX (Cgs_WT_), EMD:XXXX (Cgs_iGH1_), EMD:XXXX (Cgs_iGH2_), EMD:XXXX (Cgs_iGH3_) EMD:XXXX (Cgs_iGT_).

---

**Extended Data Fig. 1.**
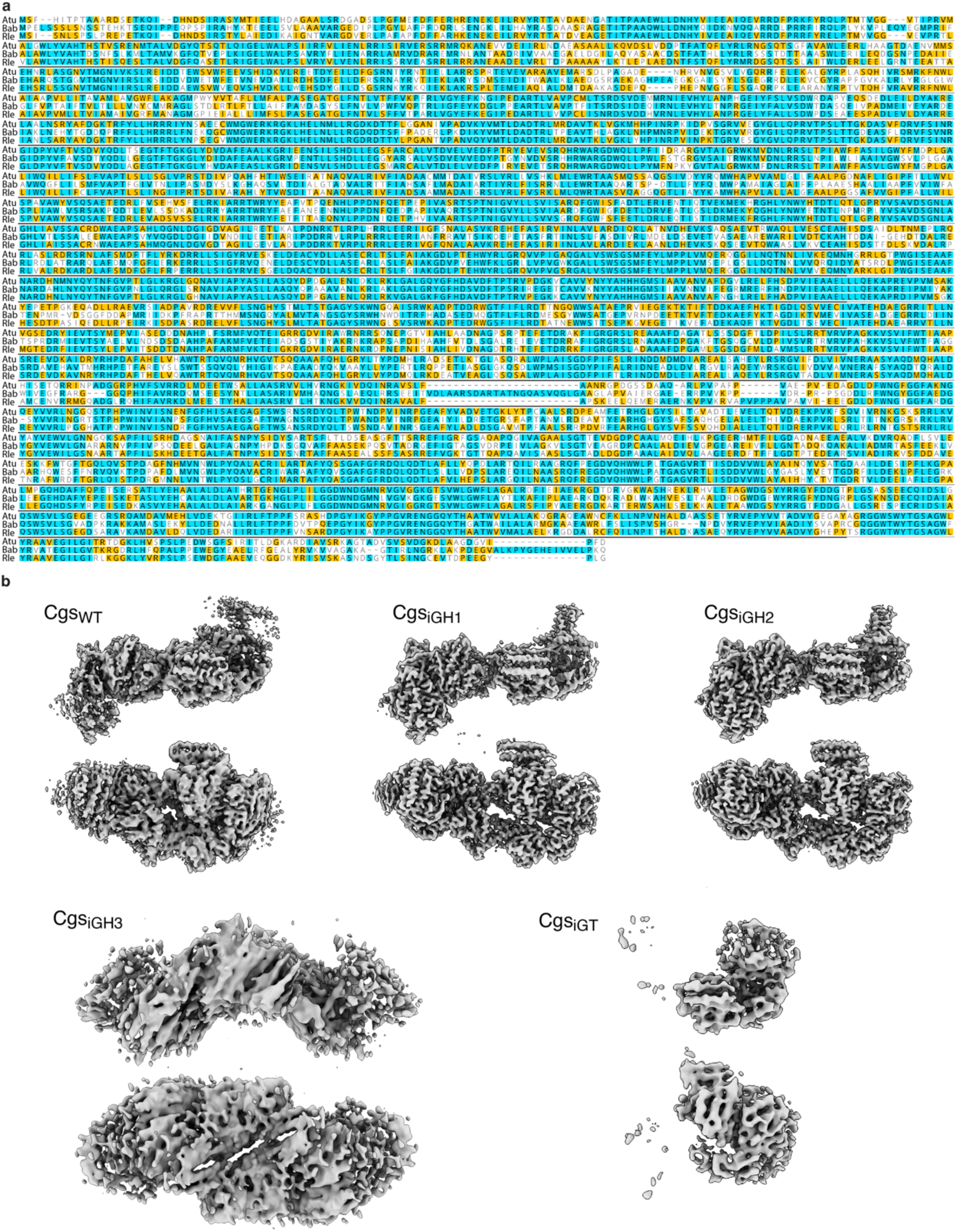
Analysis of Cgs. **a,** Sequence alignment of Cgs (NdvB) from *Agrobacterium tumefaciens* (*Atu*), *Brucella abortus* (*Bab*) and *Rhizobium leguminosarum* (*Rle*), indicating the conservation of the protein across *Rhizobiales*. **b**, Cryo-EM density maps obtained in this study. The maps were generated using the wild-type (Cgs_WT_), inactive GH94 (Cgs_iGH1_, Cgs_iGH2_, Cgs_iGH3_) and inactive GT84 (Cgs_iGT_) protein constructs. Cgs_WT_, Cgs_iGH1_ and Cgs_iGH2_ are high-resolution maps of the full-length protein. Cgs_iGH3_ represents a homodimer. Cgs_iGT_ is a low-resolution map of the N-terminal half of the protein.

**Extended Data Fig. 2.**
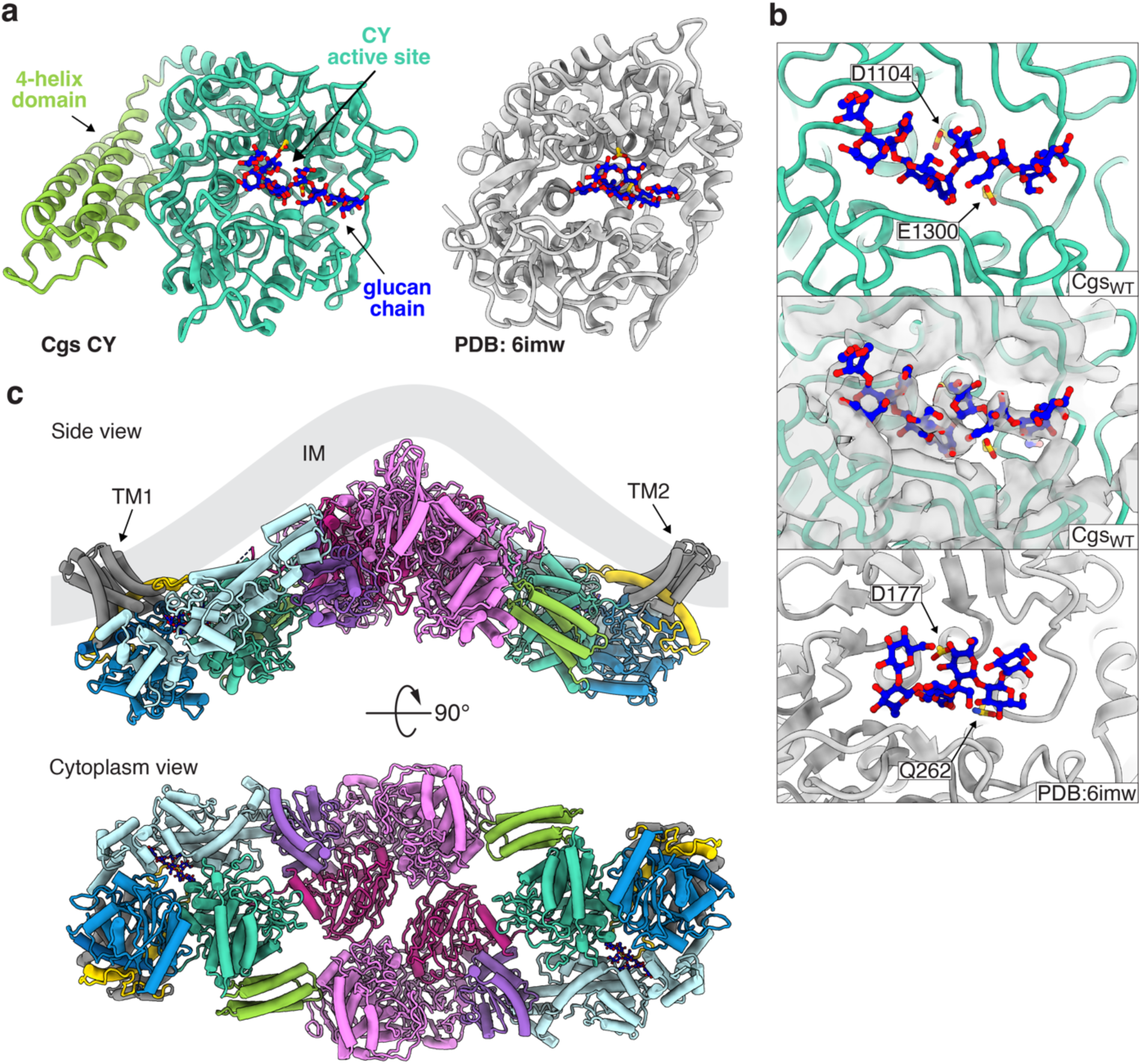
Characterization of the CY domain and its interactions. **a,** Comparison of the CY domain of Cgs (left) and the soluble GH144 homolog from *T. funiculosus* indicates a similar fold. The pairs of catalytic residues (cyan) and glucan substrates bound to the active sites (blue-red sticks) are indicated. The 120-residue part constituting the 4-helix domain (red) is absent in the homolog. **b,** Comparison of active sites of CY and GH144 models with bound substrates. Cgs CY (top, middle), as well as the *T. funiculosus* GH144 structure (PDB:6imw, bottom) are shown. Middle panel contains the density map of the glucan chain bound to CY. Active site residues are indicated in cyan. **c,** Side (top) and cytoplasm (bottom) views of the Cgs dimer structure. Cgs forms a back-to-back dimer through the interaction of the 4-helix motif with the back of the CTD. The angle between the TMs of both monomers indicates that inner membrane (IM) curvature is required for the interaction to take place.

**Extended Data Fig. 3.**
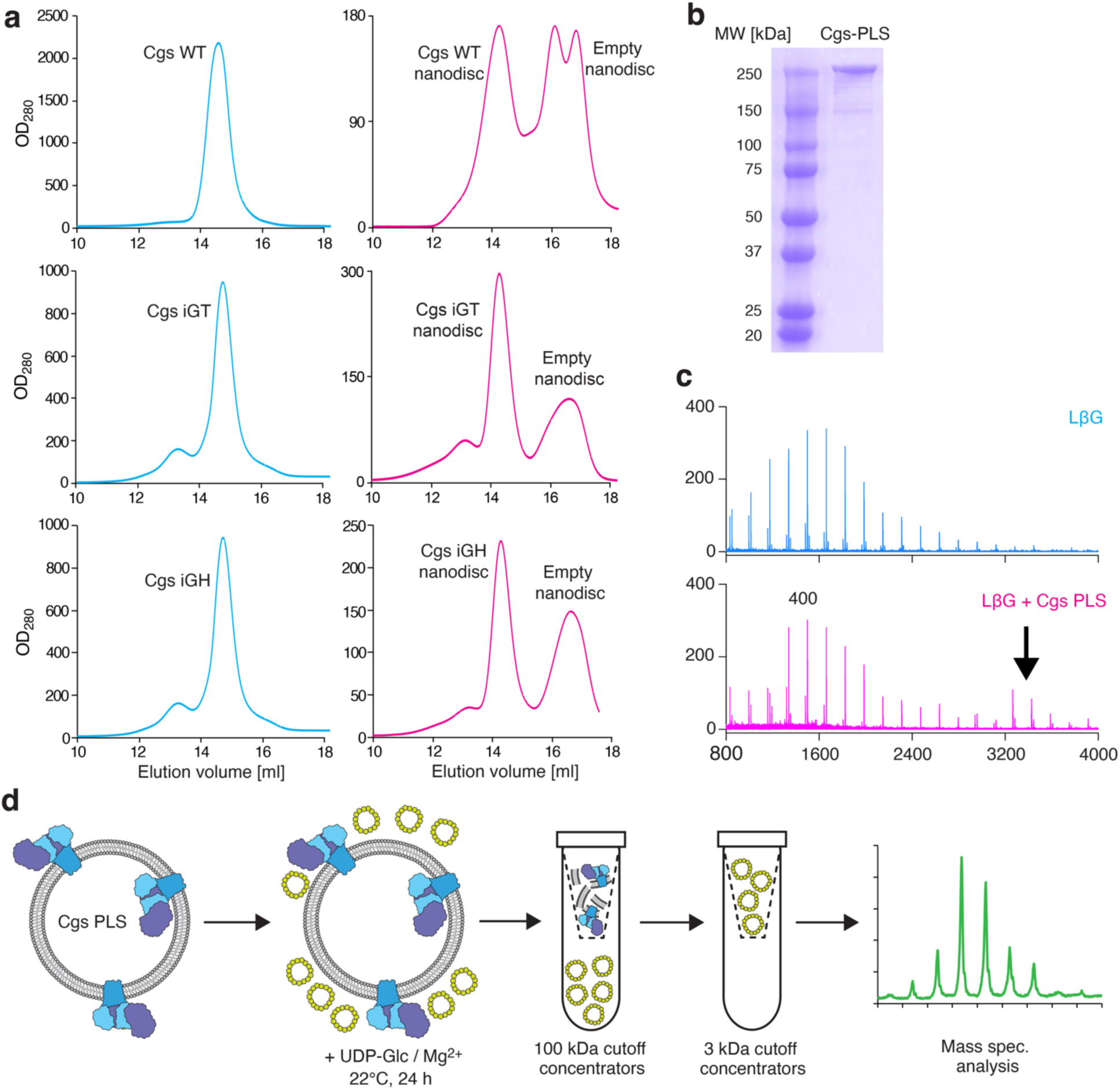
In vitro activity assay. **a,** Gel-filtration profile (Superose 6 Increase 10/300) of purified Cgs_WT_ in DDM micelles (left) and MSP1D1 nanodiscs (right). (top) Cgs_WT_; (middle) Cgs_iGT_; (bottom) Cgs_iGH_. **b,** SDS-PAGE of Cgs_WT_ proteoliposome sample indicating the incorporation of Flag-tagged Cgs (320 kDa). **c,** Processing of linear β-1,2-glucans by Cgs proteoliposomes. (top) Initial size distribution of linear β-glucans. (bottom) Size distribution of the glucan mixture after 24h incubation with Cgs proteoliposomes. **d,** Diagram describing the in vitro CβG synthesis assay with Cgs proteoliposomes.

**Extended Data Fig. 4.**
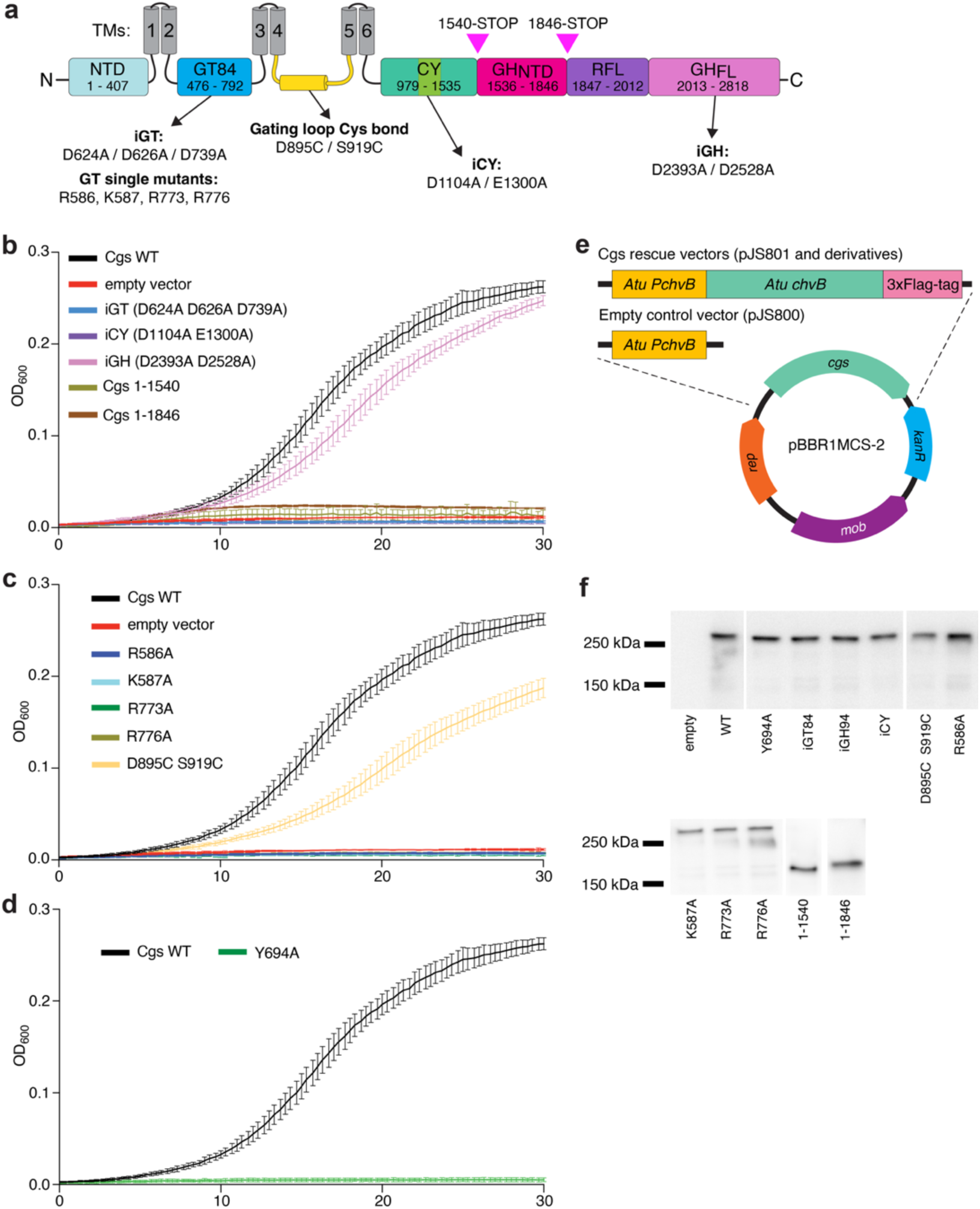
Hypoosmotic growth assay. **a,** Diagram of Cgs topology indicating the locations of mutants discussed in the study. Location of stop codons that resulted in two truncation mutants is indicated with arrowheads. **b-d,** Growth dynamic of *Atu* under osmotic stress. The *Δcgs* mutant was rescued with different constructs of the *cgs* gene. **b,** Importance of Cgs domains for function. iGH, inactive GH94 domain mutant; iGT, inactive GT84 domain mutant; iCY, inactive CY domain mutant. **c,** Role of different residues of the GT domain for function. **d,** Y694 glycosylation site is required for Cgs function. **e**, Western blot analysis of *cgs-3xFlag* gene expression in *Atu* strains used in the assay. **f**, Cartoon representation of the constructs used in the hypoosmotic stress assay. All growth curves represent mean values of 3 independent experiments (n=3). Error bars represent standard deviation. Data for graphs a-c and uncropped images for e are available as source data. Western blot analysis was performed once (n=1).

**Extended Data Fig. 5.**
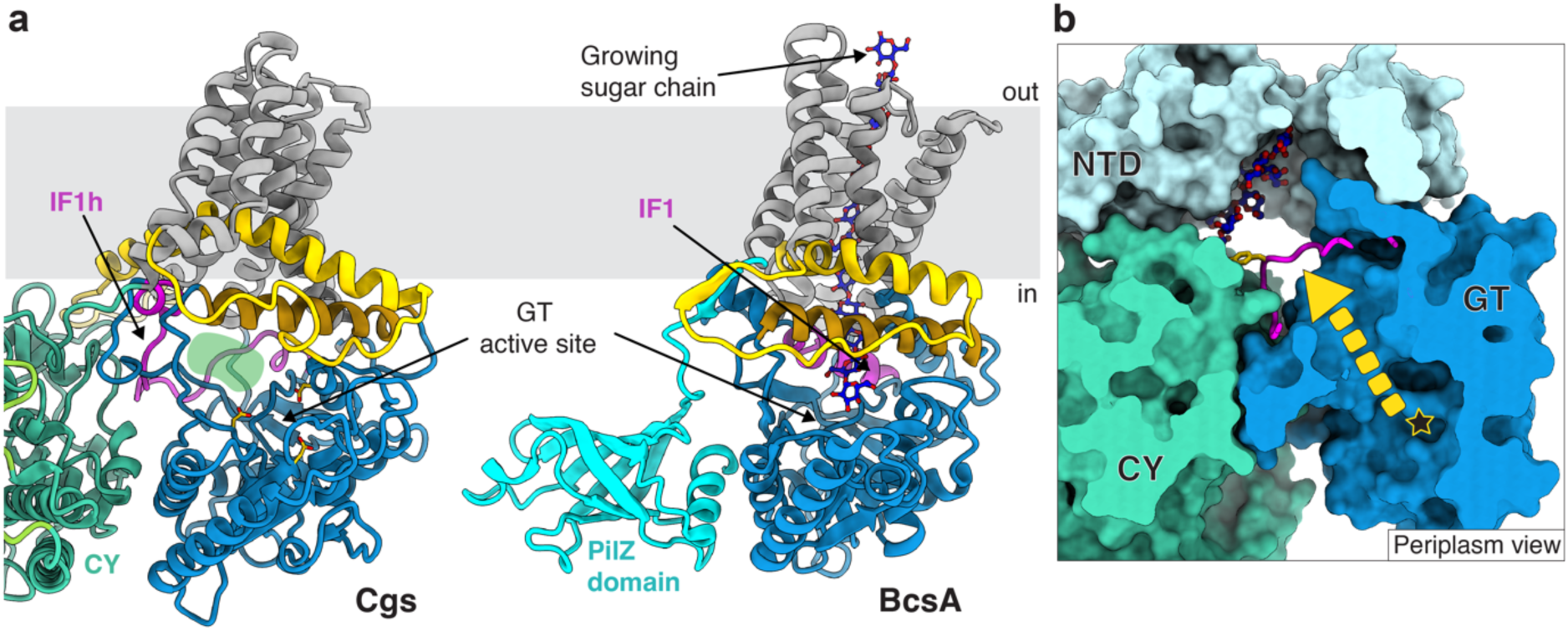
Comparison of Cgs GT84 domain with GT2 family glycosyltransferases. **a,** Comparison of Cgs (left) and BcsA (right) glycosyltransferase domains. The PilZ domain (cyan), responsible for cdGMP-dependent regulation of BcsA, is absent in Cgs. The interface motifs IF1 (magenta), IF2 (brown) and IF3 (yellow) are indicated. In BcsA, the synthesized sugar chain is translocated across a channel formed by the TM domain. The compact TM of Cgs does not form a channel. Instead, the unfolded IF1h loop opens a channel towards the other side of the GT domain (green shading). **b,** Cross-section of the channel leading from the active site of the GT (star) towards the NTD, shown from the periplasmic side. The arrow indicates the suggested path of the growing glucan chain.

**Extended Data Fig. 6.**
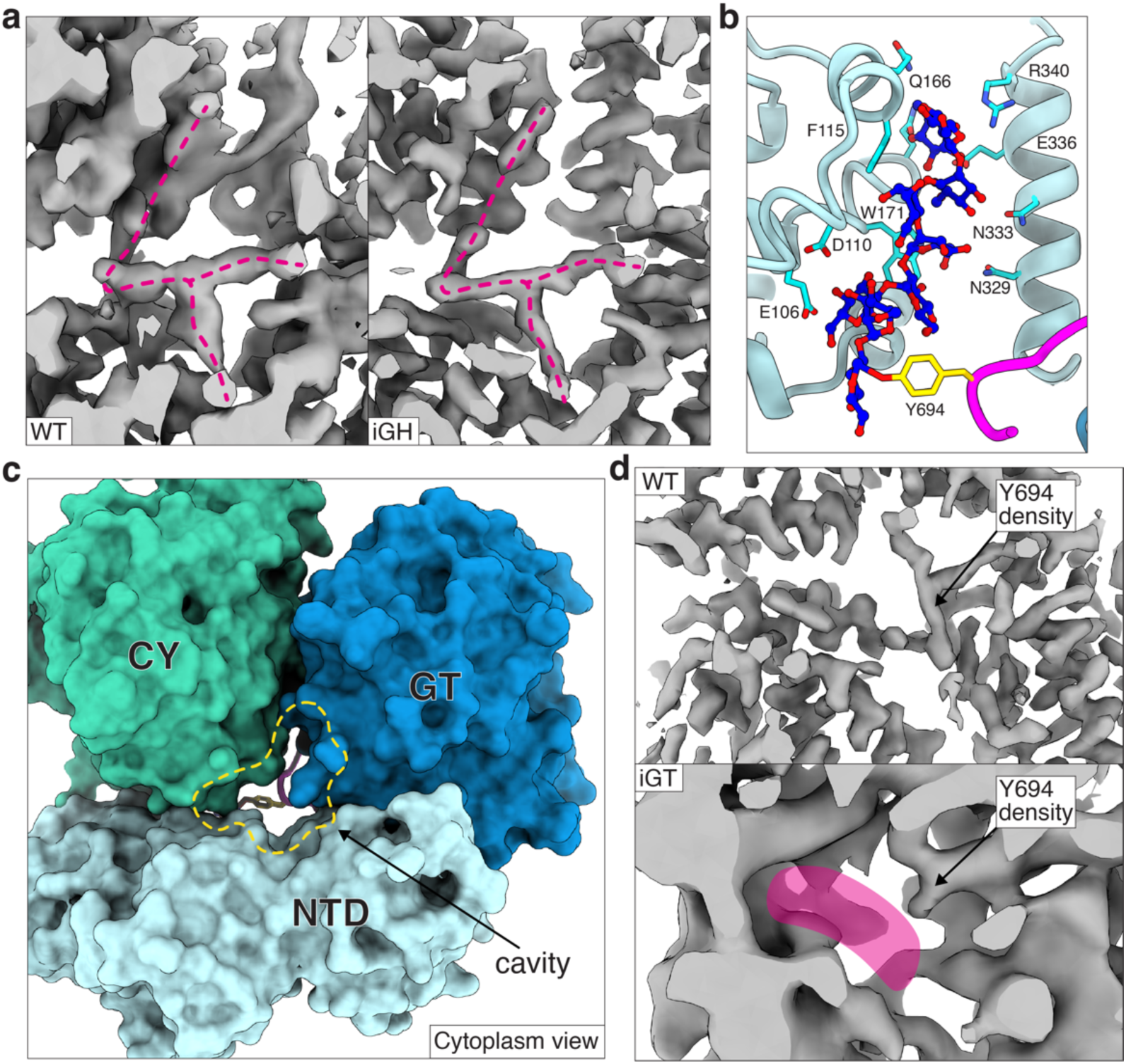
Autoglycosylation of Cgs. **a,** Cryo-EM maps obtained for the WT (left) and iGH (right) constructs. The magenta line indicates the Y-junction corresponding to the glycosylated IF1h loop. **b,** Close-up of the O-glycan (blue-red sticks) coordinated in the NTD TPR-homology fold. Residues interacting with the glucan chain are indicated in cyan. **c,** Close-up of the NTD/GT/CY interface seen from the cytoplasmic side. A large cavity (yellow line) allows access to the primer chamber from the cytoplasm. **d,** Close up of the O-glycosylation site in the iGH (top) and iGT (bottom) maps. In the low-resolution iGT structure, the volume occupied by the glucan chain appears empty (magenta).

**Extended Data Fig. 7.**
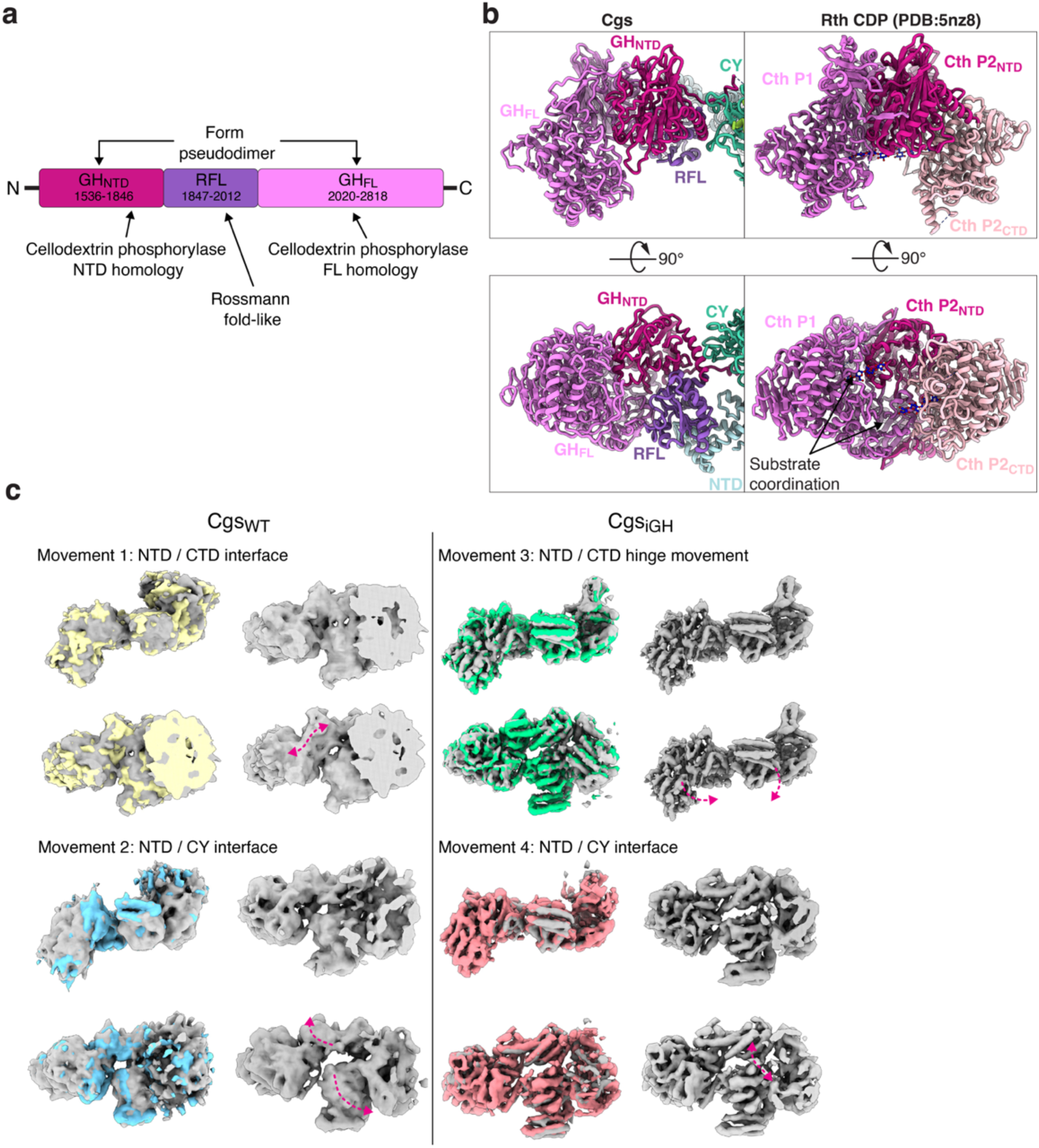
C-terminal half of Cgs. **a,** Subdomain organization of the Cgs CTD. Both ends of the CTD are homologous to the cellodextrin phosphorylase-domain from the GH94 family: a shorter, NTD fragment at the N-terminus (GH_NTD_) and a full-length subdomain on the C-terminus (GH_FL_). They are separated by a small Rossmann fold-like subdomain (RFL). **b,** Comparison of the Cgs CTD (left) with the crystal structure of a soluble homolog CDP (PDB:5nz8, right) from *C. thermocellum* (Cth). The GH_NTD_ and GH_FL_ form a pseudodimer that reflects the dimerization of CDP. The cellotetraose substrate bound to the CDP dimer indicates that the dimerization interface plays a role in coordinating the substrate chain into the active site of the phosphorylase (bottom right). The CTD of the second CDP molecule (P2_CTD_) that is not present in Cgs is colored in pale pink. **c,** 3D variability analysis of Cgs_WT_ (left) and Cgs_iGH_ (right) maps. Large domain movements are indicated with arrows. Relative domain movement leads to changing distance at the interfaces between NTD and CTD (1), as well as NTD and CY (2, 4). A hinge movement between the N- and C-terminal halves of Cgs can be observed (3). The opening between the NTD and CY in some conformations allows access from the primer chamber to the CY active site.

**Extended Data Fig. 8.**
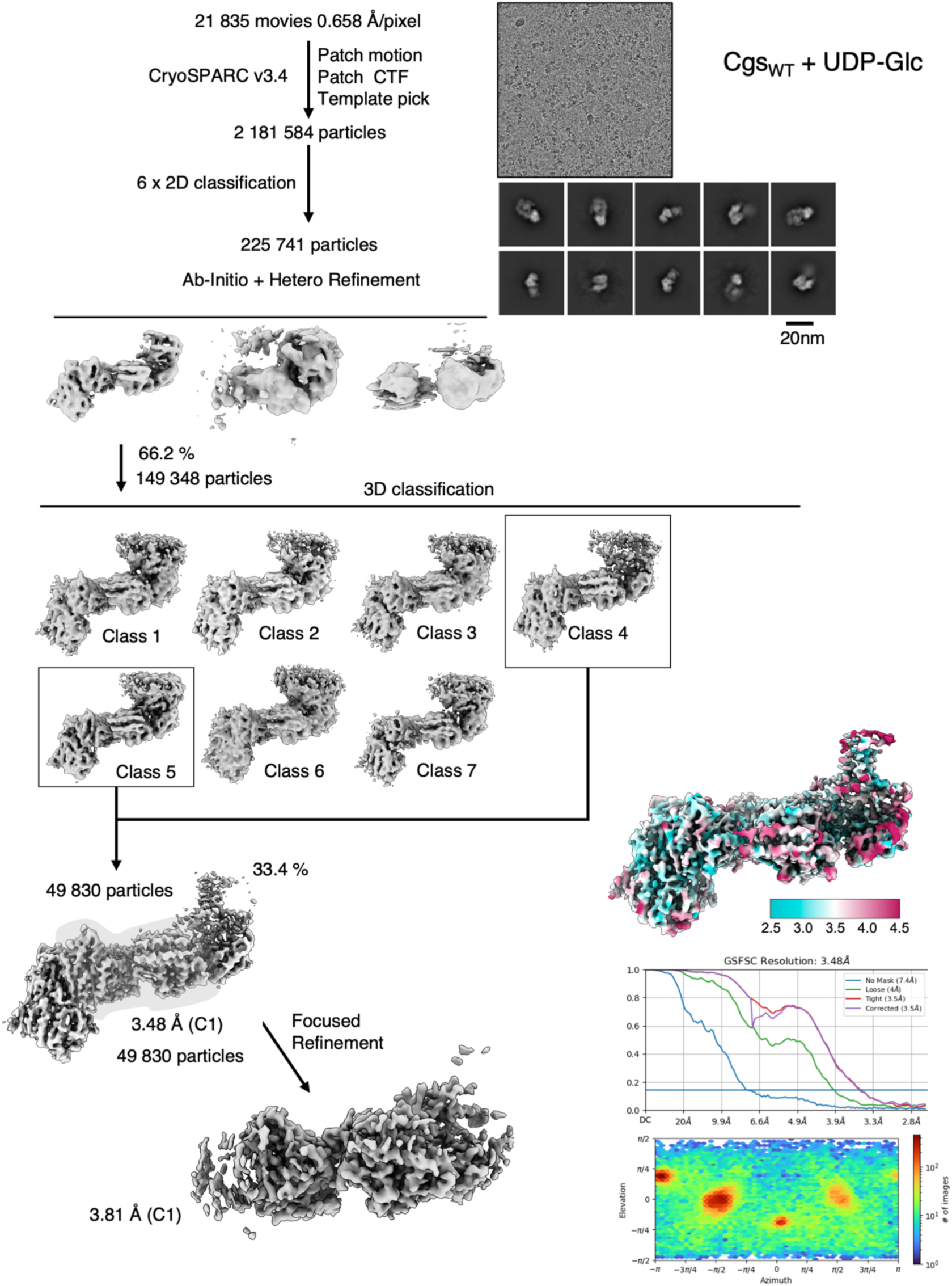
Single particle processing workflow of the dataset obtained with the Cgs_WT_ UDP-Glc sample.

**Extended Data Fig. 9.**
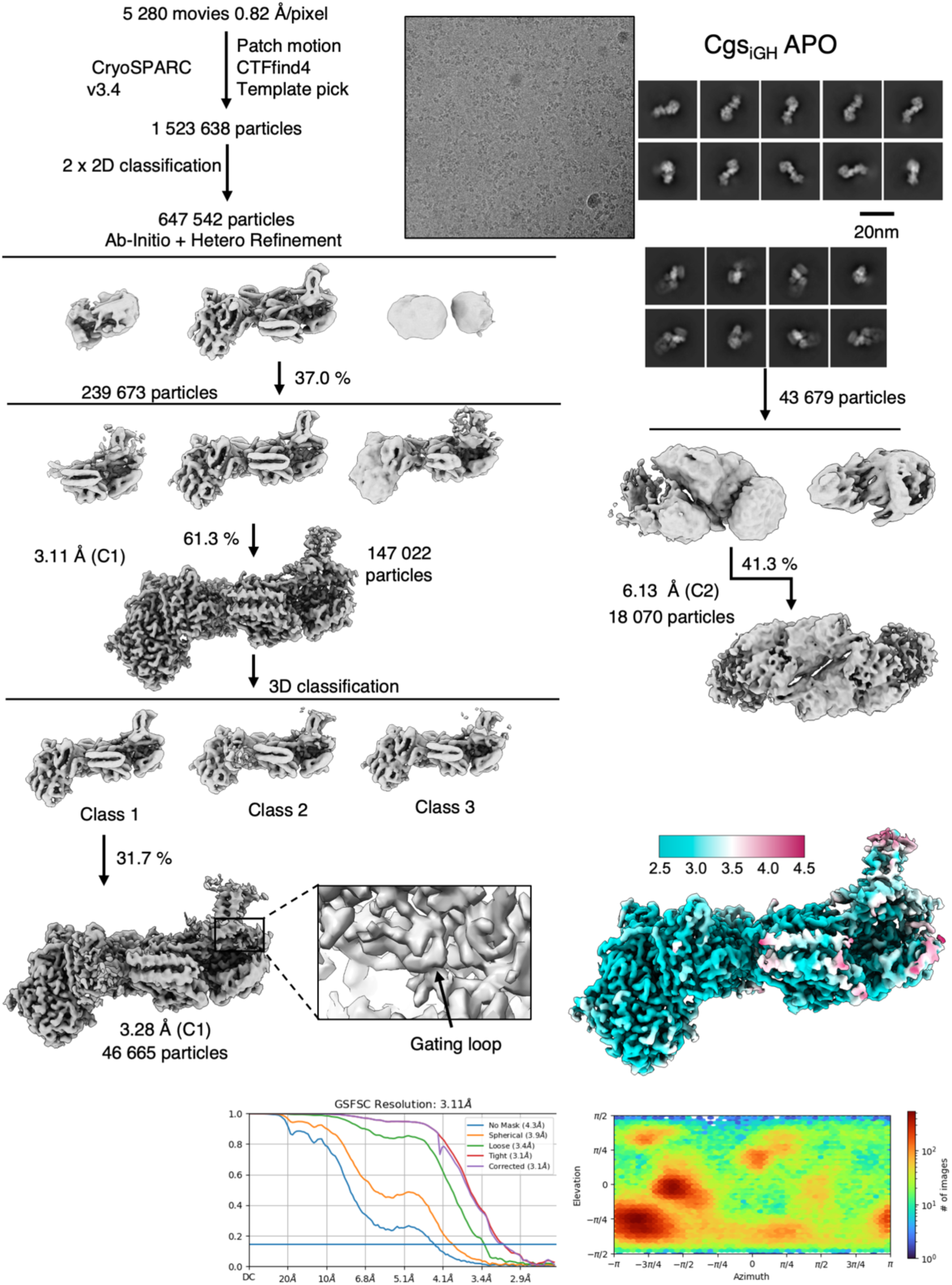
Single particle processing workflow of the dataset obtained with the Cgs_iGH_ APO sample.

**Extended Data Fig. 10.**
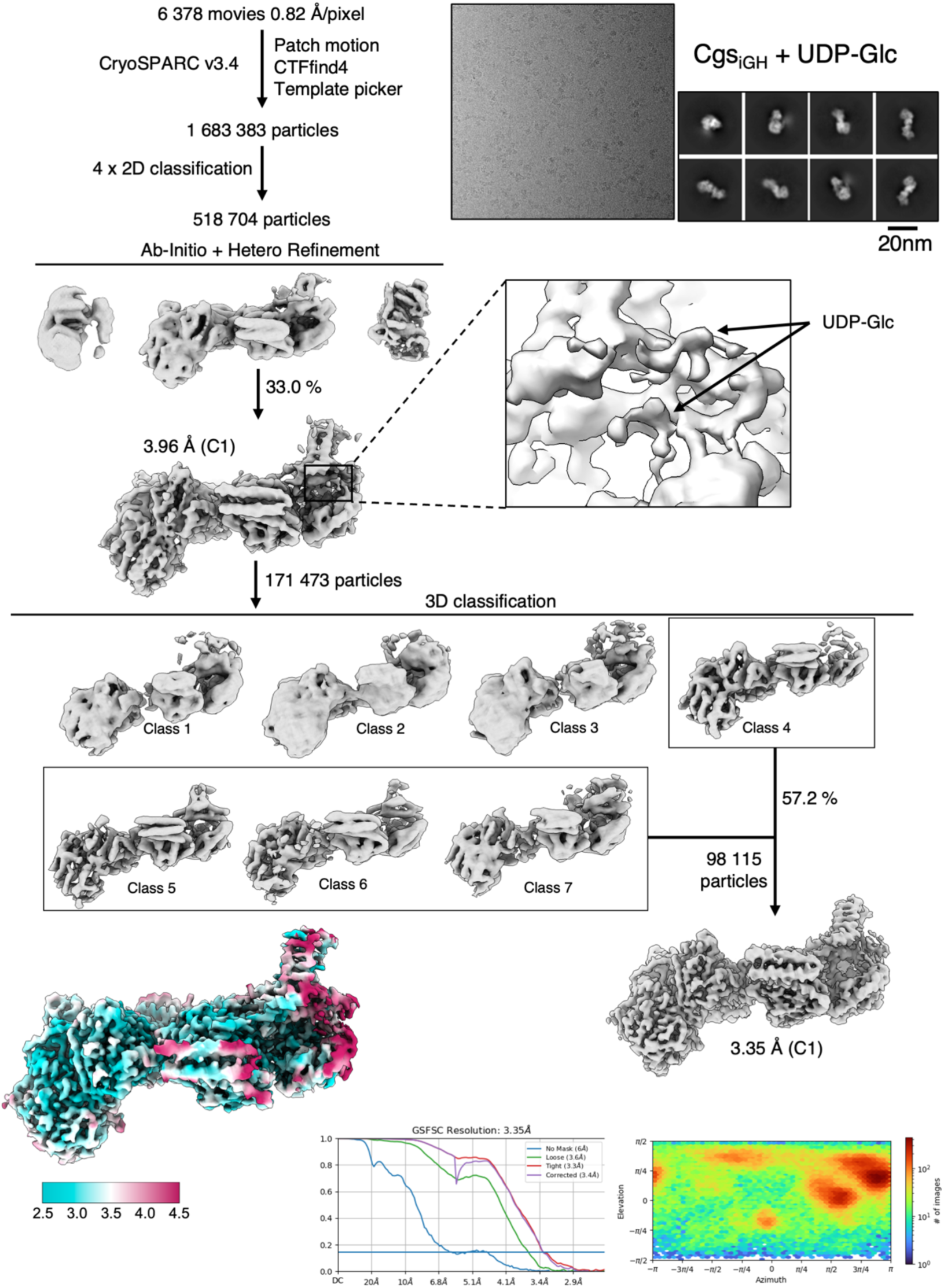
Single particle processing workflow of the dataset obtained with the Cgs_iGH_ UDP-Glc sample.

**Supplementary Table 1.**
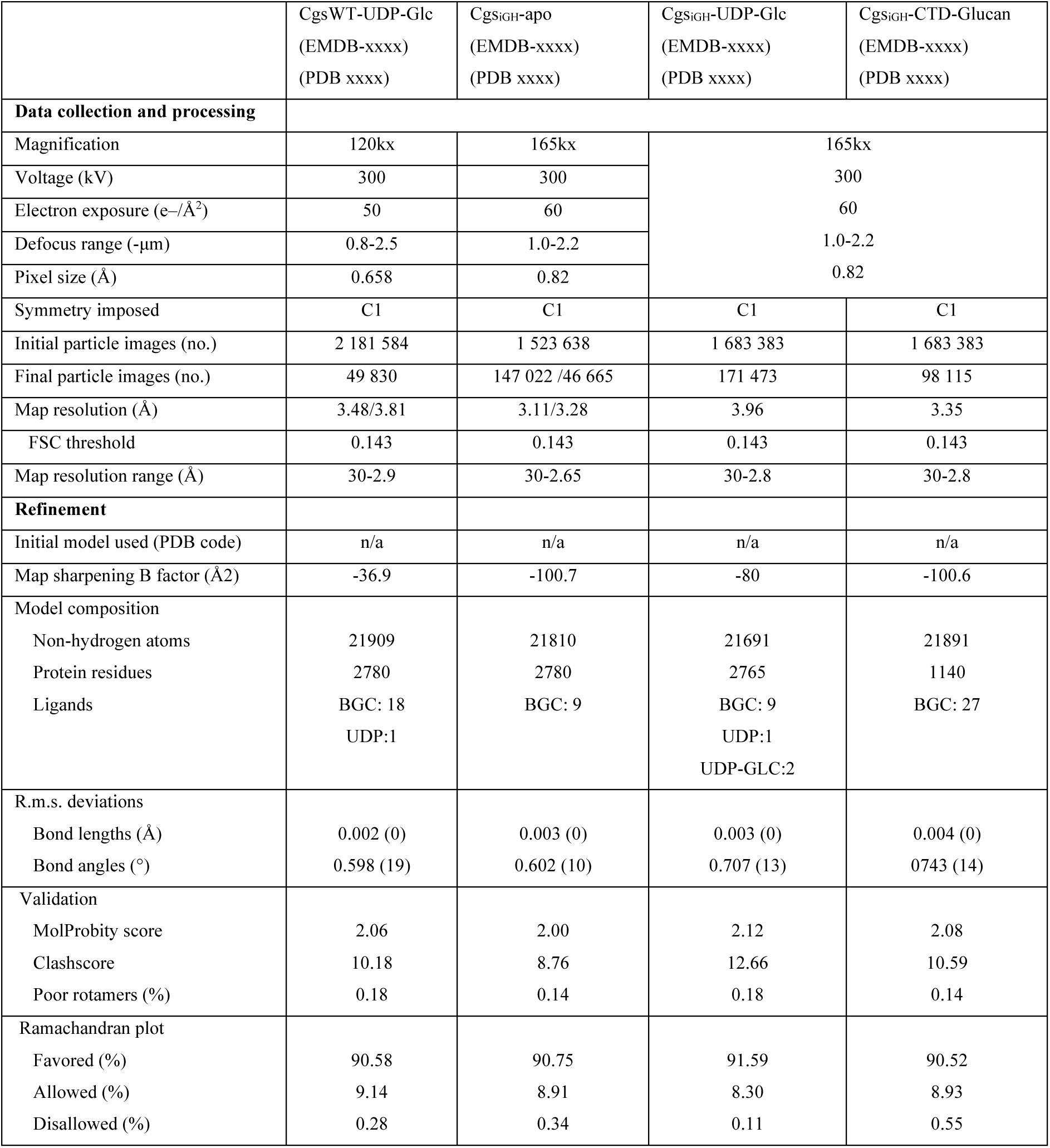
Cryo-EM data collection, refinement and validation statistics.

**Supplementary Table 2.**
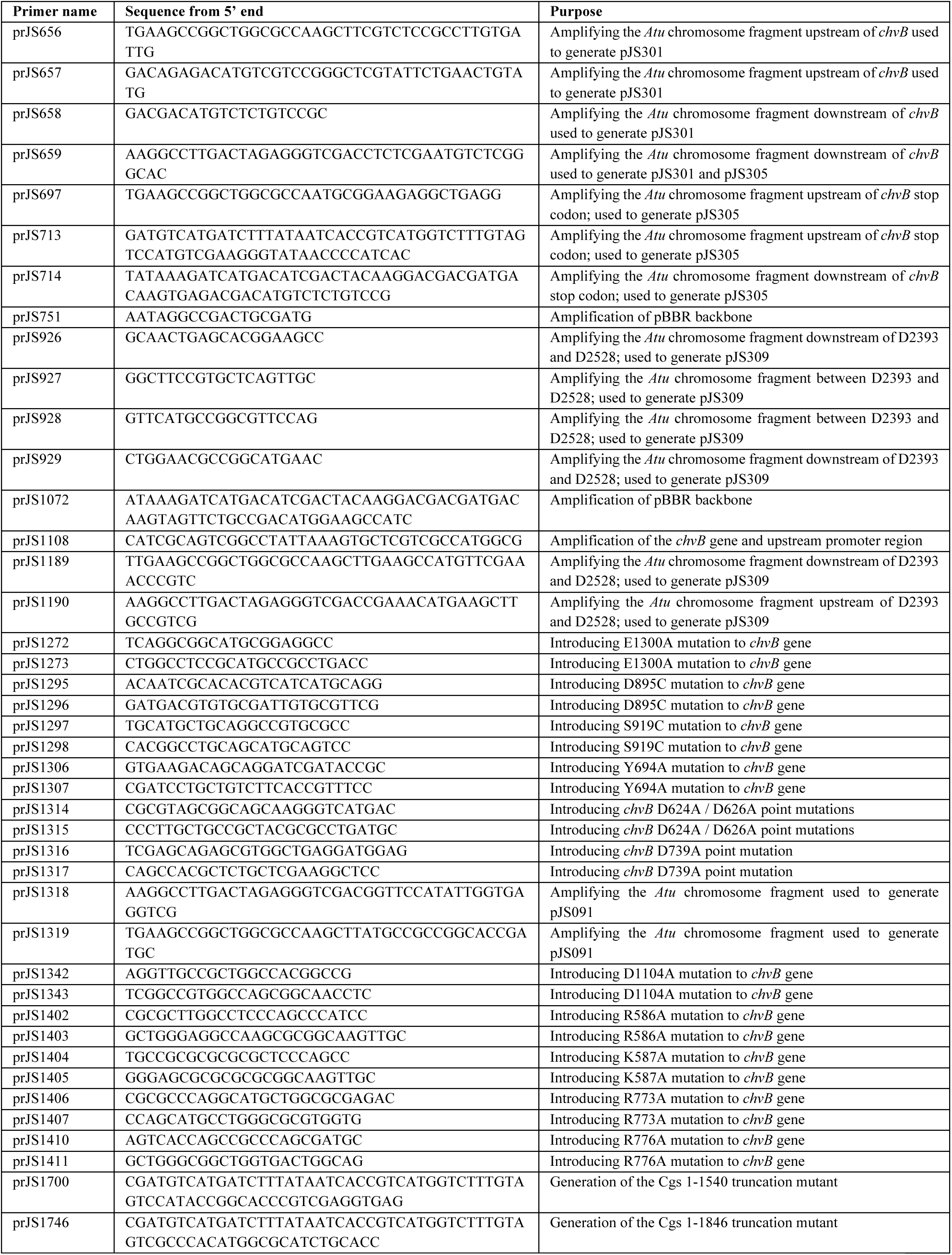
Primers used in this study.

**Supplementary Table 3.** Plasmids generated for this study.

| Plasmid name | Plasmid background | Description |
| --- | --- | --- |
| pJS091 | pNPTD138 | Introduces D624A/D626A/D739A mutations to the genomic *chvB* gene |
| pJS301 | pNPTD138 | Full length deletion of *chvB* gene |
| pJS305 | pNPTD138 | Adds a genomic C-terminal 3xFlag-tag to *chvB* gene |
| pJS309 | pNPTD138 | Introducing D2393A/D2528A mutations to the genomic *chvB* gene |
| pJS800 | pBBR-MCS1 | Empty control vector |
| pJS801 | pBBR-MCS1 | Rescue vector – cgs WT |
| pJS802 | pBBR-MCS1 | Rescue vector – cgs Y694A |
| pJS803 | pBBR-MCS1 | Rescue vector – cgs iGT84 (D624A/D626A/D739A) |
| pJS804 | pBBR-MCS1 | Rescue vector – cgs iCY (D1104A/E1300A) |
| pJS806 | pBBR-MCS1 | Rescue vector – cgs iGH94 (D2393A/D2528A) |
| pJS820 | pBBR-MCS1 | Rescue vector – cgs R586A |
| pJS821 | pBBR-MCS1 | Rescue vector – cgs K587A |
| pJS822 | pBBR-MCS1 | Rescue vector – cgs R773A |
| pJS823 | pBBR-MCS1 | Rescue vector – cgs R776A |
| pJS829 | pBBR-MCS1 | Rescue vector – cgs D895C S919C |
| pJS862 | pBBR-MCS1 | Rescue vector – cgs 1-1540 truncation mutant |
| pJS865 | pBBR-MCS1 | Rescue vector – cgs 1-1846 truncation mutant |

**Supplementary Table 4.** *Agrobacterium* strains used in this study.

| Strain name | Carried plasmid | Description |
| --- | --- | --- |
| Atu009 | - | *A. tumefaciens* C58 with 3xFlag-tag insertion before the stop codon of *chvB* |
| Atu004 | - | *A. tumefaciens* C58 with full length deletion of *chvB* gene |
| Atu125 | - | Atu009 with additional D2393A/D2528A mutations in *chvB* |
| Atu130 | - | Atu009 with additional D624A/D626A/D739A mutations in *chvB* |
| Atu243 | pJS800 | Atu004 with empty control vector |
| Atu219 | pJS801 | Atu004 with rescue plasmid encoding for Cgs WT |
| Atu220 | pJS802 | Atu004 with rescue plasmid encoding for Cgs Y694A |
| Atu221 | pJS803 | Atu004 with rescue plasmid encoding for Cgs iGT (D624A/D626A/D739A) |
| Atu222 | pJS804 | Atu004 with rescue plasmid encoding for Cgs iCY (D1104A/E1300A) |
| Atu224 | pJS806 | Atu004 with rescue plasmid encoding for Cgs iGH (D2393A/D2528A) |
| Atu230 | pJS820 | Atu004 with rescue plasmid encoding for Cgs R586A |
| Atu231 | pJS821 | Atu004 with rescue plasmid encoding for Cgs K587A |
| Atu232 | pJS822 | Atu004 with rescue plasmid encoding for Cgs R773A |
| Atu233 | pJS823 | Atu004 with rescue plasmid encoding for Cgs R776A |
| Atu239 | pJS829 | Atu004 with rescue plasmid encoding for Cgs D895C S919C |
| Atu260 | pJS862 | Atu004 with rescue plasmid encoding for Cgs 1-1540 truncation mutant |
| Atu355 | pJS865 | Atu004 with rescue plasmid encoding for Cgs 1-1846 truncation mutant |

